# Tissue-autonomous phenylpropanoid production is essential for establishment of root barriers

**DOI:** 10.1101/2020.06.18.159475

**Authors:** Tonni Grube Andersen, David Molina, Joachim Kilian, Rochus Franke, Laura Ragni, Niko Geldner

**Affiliations:** Max Planck Institute for Plant Breeding Research, Carl-Von-Linné-weg 10, 50829, Cologne, Germany; ZMBP-Center for Plant Molecular Biology, University of Tuebingen, Auf der Morgenstelle 32, 72076 Tuebingen, Germany; Institute of Cellular and Molecular Botany, Rheinische Friedrich-Wilhelms-University of Bonn, Kirschallee 1, 53115, Bonn, Germany; Department of Plant Molecular Biology, University of Lausanne, 1015 Lausanne, Switzerland

## Abstract

Plants deposit polymeric barriers in their root cell walls to protect against external stress and facilitate selective nutrient uptake. The compounds that make up these barriers originate from the fatty acid- and phenylpropanoid biosynthetic pathways. Although the machinery responsible for production of the barrier constituents is well-characterized, our pathway models lack spatiotemporal resolution – especially in roots - and the source tissue is often not clear due to the apoplastic nature of barriers. Insights into how the individual root tissues or cells contribute to forming apoplastic barriers is important for elucidation of their ultrastructure, function and development. Manipulation of the associated biosynthesis is delicate, as mutants often display pleiotropic phenotypes due to the broad role of the underlying metabolites. Here, we address these issues by creating a genetic tool that allows in vivo repression of the phenylpropanoid pathway with both spatial and temporal control. We provide strong evidence that tissue-autonomous production of phenylpropanoids is essential for establishment of the endodermal Casparian strip. Moreover, we find that in order to maintain deposition and attachment of a coherent suberin matrix to the cell wall, cells require continuous production of aromatic constituents. This process is especially crucial in the suberized endodermis where we find that repression of phenylpropanoid production leads to active removal of suberin.

## INTRODUCTION

Among the diffusion barriers deposited in root cell walls, the best characterized is, arguably, the Casparian strip (CS), which is initiated between the newly differentiated endodermal cells. The CS consists of precisely localized oxidatively-coupled lignin-polymer depositions in the apoplastic space^1,2^. As the endodermal development progresses, the entire surface of most cells is covered by hydrophobic suberin depositions^3^. In contrast to the CS, the suberin-barrier is a lamellae-like structure deposited below the primary cell wall^4,5^, serving as a transmembrane rather than an apoplastic barrier^6–8^. These suberin depositions consist of a crosslinked matrix of aliphatic fatty acids, glycerol and aromatic monomers joined through a variety of ester-formations and oxidative couplings^9,10^. Establishment and function of endodermal barriers is tightly controlled by a surveillance system^11^. This system consists of an elegant, localized multi-component pathway^12^, which requires diffusion of Casparian strip Integrity Factor (CIF) peptides from the stele to the cortex-facing surface of the endodermis. Here, CIF peptides bind and activate the Leucine Rich Repeat (LRR)-family receptor SCHENGEN 3^13^ (SGN3, also called GSO1^14^). This creates a self-regulating system, as formation of a tight CS inhibits CIF diffusion. Importantly, in the case of a non-functional CS, excess activation of this pathway serve as a signal to initiate endodermal “sealing” through ectopic non-CS localized lignification and suberin deposition in the young, normally unsuberized endodermis^15^.

During secondary growth, the root increases in girth, the endodermis undergoes programmed cell death and its barrier function is replaced by the periderm^16^. The periderm is a dynamic structure comprising the meristematic phellogen, which divides bifacially and gives rise to cork (toward the soil) and phelloderm (towards the vasculature) layers. The cork cell walls are highly suberized and lignified, which confers the barrier property to the periderm. Moreover, elevated number of cork layers and suberin content have been associated with increased tolerance to certain stresses^16–19^.

The metabolites required for root barrier formation are derived from the fatty acid (FA, aliphatic) and the phenylpropanoid (PP, aromatic) pathways. For production of aliphatic suberin constituents, FAs are elongated, oxidized^20,21^ and conjugated to glycerol via glycerol-3-phosphate acyltransferases (GPATs)^22–24^. Suberin biosynthetic gene expression correlates well with suberin deposition^25,26^ and *GPAT5* has been established as a marker for endodermal cells undergoing suberization^27^. Thus, it is clear that, in roots, aliphatic constituents of suberin are produced by the depositing tissues^3^. PP synthesis gives rise to distinct compounds, which serve multiple roles throughout the lifespan of the plant. Besides acting as structural constituents, such as coniferyl alcohol (lignin) and ferulic acid (suberin), PP-derived aromatic metabolites also function in chemical systems such as defenses, chelators and/or pollinator attractors^28^. Phenylalanine is the major precursor for PPs in most plants and the initial common, committed steps are catalyzed by the enzymes Phenylalanine Ammonia Lyase (PAL)^29^ and Cinnamate-4-Hydroxylase (C4H)^30–32^. For the barrier-forming tissues, the precursors needed for polymerization and deposition in the cell walls must be transported across the plasma membrane to reach the apoplast. As a consequence, integrated metabolites may be produced and employed cell-autonomously or originate from neighboring or distal cells. For suberin, the tight connection of FA- and PP-derived metabolites required, render diffusion from surrounding tissues unfeasible, which is in line with our current models of tissue-autonomous suberin production. Yet, in the xylem, both cell-autonomous and distal PP production for lignin monomers occurs^33^. Thus, for the lignin-based CS, the endodermis probably rely on diffusion from the underlying xylem, or be an autonomous producer of the required compounds.

In this work, we address the ability of the endodermis and periderm of Arabidopsis roots to synthesize PP-derived metabolites. We show that essential PP biosynthesis genes are expressed in both endodermis and cork. By employing transcriptional repressors, we create plants with controllable reduction in PP synthesis within discrete, tissue-specific zones. This allowed us to investigate the origin of PP precursors and the functional role in different root barriers across plant development. We found that continuous tissue-autonomous production of PP-derived compounds within the zone of CS establishment is essential for formation of this barrier. Our data additionally highlight that in contrast to the cork barrier, which appears to represent an end-point decision for suberin deposition, endodermal suberin is dynamic and stereotypical endodermal suberin lamellae likely represent a steady-state between continuous synthesis and degradation.

## RESULTS

### Endodermal cells have the capacity for phenylpropanoid synthesis

Arabidopsis has 4 homologues of the *PAL* genes^34^, where only *PAL1,2* and *4* transcripts have been detected in roots^35^ (Supplementary Figure 1). In contrast, only one C4H locus exists^34^, with strong ubiquitous expression in all root tissues^35^ (Supplementary Figure 1). To increase the resolution of these patterns, we constructed fluorescent, transcriptional reporter lines based on promoter regions of *PAL1,2,4* and *C4H* driving expression of a nuclear-localized triple mVenus reporter (NLS 3xmVenus). In 5-day-old seedlings, activity of all promoters could be observed within the root vasculature (Figure 1), which is expected to be related to synthesis of lignin for xylem formation. However, *PAL1* and *2* promoter activity was additionally observed in the early stages of endodermis adjacent to the lignifying protoxylem (Figure 1). *PAL4* activity was also found in the endodermis, yet this appeared to be specifically in older, possibly suberizing, cells (Figure 1). In addition to a similar strong activity in the vasculature, the *C4H* promoter was confirmed to be expressed across all investigated stages of the endodermal development (Figure 1). Thus, the endodermis appears to express key genes for PP production, making it likely that this layer has capacity for PP synthesis.

**Figure 1.**
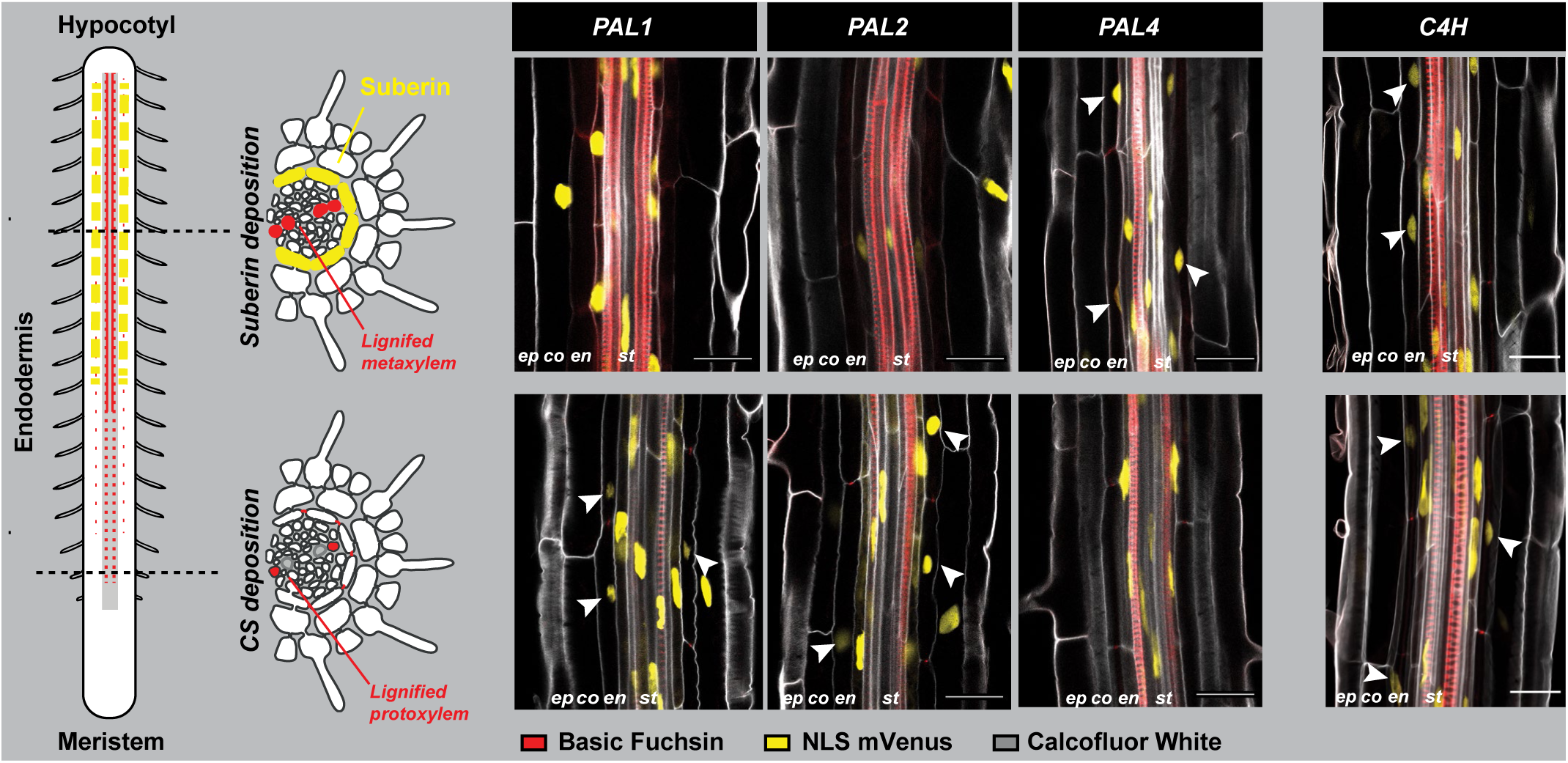
Essential phenylpropanoid pathway genes are expressed in the root endodermis. Activity of nuclear localized (NLS) 3x mVenus fusion reporters driven by promoter regions of root-expressed phenyl alanine lyase (*PAL*) and cinnamate 4-hydroxylase (*C4H*) homologues in Arabidopsis roots. Expression in in 5-day-old roots in zones of lignification (bottom) and suberization (top) of the endodermis. All roots were fixed using a previously established ClearSee protocol^54^. Cell walls were highlighted using Calcofluor White, whereas Basic Fuchsin was used to highlight lignin depositions in the xylem and endodermis. Arrowheads point to endodermis cells with reporter activity. co: Cortex, en: Endodermis ep: Epidermis, st: Stele. Scalebars represent 25 *μ*m.

### Lignification of CS and xylem follows similar recovery dynamics

To investigate if the lignin deposition in the CS and xylem is coordinated, we set out to study the onset of lignification after disruption of PP synthesis in the entire root. By treating 5-day-old Arabidopsis seedlings for 24 hours with the C4H-inhibitory chemical piperonylic acid (PA)^36^, we could create non-lignified zones where the developing xylem and CS would normally have initiated lignification^37^. We then transferred seedlings to recovery plates (without PA), in presence or absence of the lignin monomer coniferyl alcohol. This allowed us to independently address the capacity of the apoplastic lignin polymerization machinery and the recovery of endogenous lignin monomer production (Figure 2a). We measured recovery of lignification over a span of several hours and observed a significant (P<0,01, Students T-test vs untreated) lignin-specific fluorescence (Basic Fuchsin stain) for both the xylem and CS in this zone after only 4 hours of transfer to recovery plates (Figure 2b, Supplementary Figure 2). This was decreased to 2 hours for both zones in presence of 20 *μ*M coniferyl alcohol (Figure 2b and Supplementary Figure 2a), which confirms that PA limits the endogenous substrate availability, but not the apoplastic polymerization capacity. Only in the presence of externally supplied coniferyl alcohol did both the xylem zone and CS recover to nearly 100% of the signal observed in control plants within the tested timeframe (Figure 2b). This probably reflects a decoupling of the polymerization capacity from substrate availability due to the recovery of C4H activity from PA inhibition. In all cases, onset of lignification in xylem and CS showed similar dynamics (Figure 2b) and appeared at similar stages of the early root (Supplementary Figure 2a), which might indicate coordination of PP synthesis in the distinct tissues.

**Figure 2.**
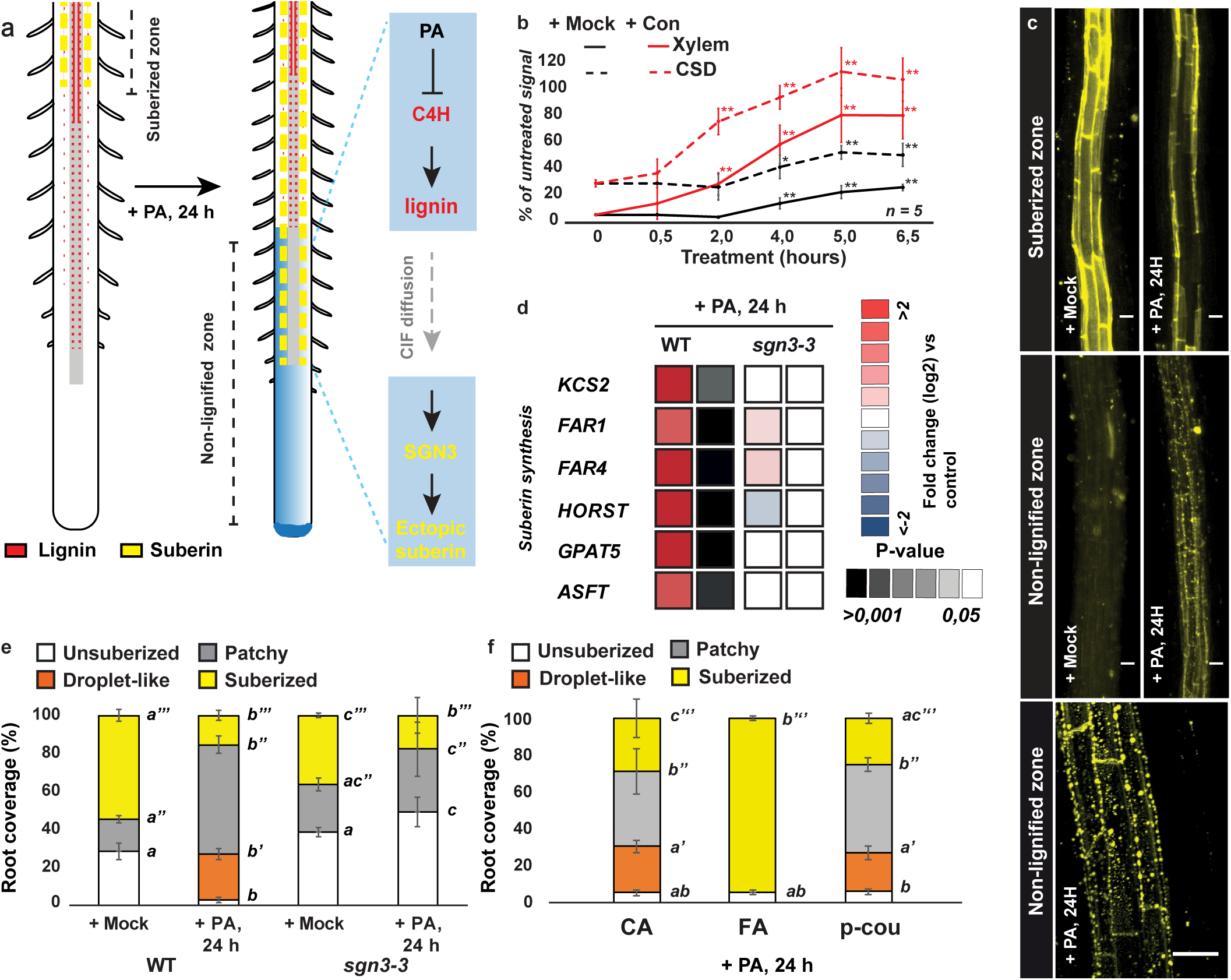
Endodermal suberin deposition is disrupted by inhibition of PP synthesis and specifically com-plemented by FA. a) Upon treatment of 4-day-old roots with the phenylpropanoid synthesis inhibitor Piperonylic acid (PA) for 24 hours, the continuous root-growth leads to establishment of a non-lignified zone where the xylem and Casparian strip (CS) are normally lignified. Absence of lignification in the CSD leads to diffusion of Casparian strip Integrity Factor peptides (CIF) and consecutive activation of the Schengen pathway^15^. This results in ectopic suberization of the otherwise non-lignified endodermal cells. b) Time course analysis of Basic Fuchsin signal recovery in the non-lignified zone of xylem or CSD after 24 h of PA pre-treatment. 6-day-old plants were moved from PA-containing plates to recovery plates with either a mock solution or 20 *μ*M coniferyl alcohol (Con) at the indicated timepoints. The signals were normalized to identically grown non-PA treated plants. N= 5, * P< 0,05; ** P<0,01 Two-sided Students T-test vs 0 hour treatment. c) Fluorol Yellow (FY) staining of suberin in the suberized and non-lignified zones of 5-day-old Arabidopsis seedlings treated with mock or PA for 24 h before staining. In the non-lignified zone of PA-treated roots, FY-staining gave rise to “droplet-like” structures as seen in the lowermost panel. d) Root-specific differential expression analysis of genes involved in suberin biosynthesis upon 24 h PA treatment of 5-day-old plants. Expression was normalized to non-PA treated control plants for each genotype. P-values are based on a two-sided Students T-test (PA-vs mock-treated). e) Quantification of FY stained endodermal suberization in 5-day-old Arabidopsis WT or *sgn3-3* seedlings treated with mock (EtOH) or PA for 24 h. N=6. f) Quantification of FY stained endodermal suberization in 5-day-old Arabidopsis WT seedlings treated with PA for 24 h in combination with mock (EtOH), 20 *μ*M caffeic (CA), ferulic (FA) or p-coumaric acid (p-cou). N=6. All error bars are SD, Letters refer to individual groups in a one-way ANOVA analysis with a post-hoc multiple group T-test (Tukey). Scale bars represent 25 *μ*m.

### Inhibition of phenylpropanoid synthesis leads to aberrant suberin deposition

As disruption of lignification in the CS promotes ectopic suberization in young parts of the endodermis due to the SGN pathway^11–13^, our experimental setup allowed us to additionally assess *de novo* deposition of endodermal suberin in a zone incapable of producing PP-derived metabolites (Figure 2a). Instead of showing a smooth FY signal typical of wild-type (WT) endodermal suberin deposition (Supplementary Figure 2b), PA-treated roots had a “droplet-like” FY staining in this non-lignified zone. Within the zone of suberization onset (patchy zone), we could observe droplet- and reticulate-like structures (Supplementary Figure 2b). We did not see this phenomenon in the *sgn3-3* mutant with a non-functional SGN pathway (Figure 2c and e, Supplementary Figure 2b), and saw a strong SGN-dependent upregulation of genes related to suberin biosynthesis after 24 h PA treatment (Figure 2d). Surprisingly, in PA-treated roots, the zone of continuous suberization in the late endodermis, already established before start of treatment, was strongly decreased in a manner that was independent of the SGN pathway (Figure 2c and e, Supplementary Figure 2b). Intriguingly, this suggests that pre-existing suberin lamellae are removed or degraded upon PP synthesis inhibition. By addition of suberin-associated PP metabolites to the plates, we found that presence of 20 *μ*M ferulic acid in the media led to complementation of the droplet-like ectopic suberin deposition and disappearance in the pre-established zone (Figure 2f), Supplementary Figure 2b). Thus, tight coordination of aromatic and aliphatic monomer synthesis is necessary for correct establishment of the suberin barrier. As this coordination is inevitably intertwined with spatial organization between tissues, we set out to investigate the role of the endodermis in PP synthesis, and to clarify if construction of the endodermal barriers is coordinated with lignin production in the xylem.

### Monolignols required for CS formation are produced in the endodermis

Transcriptional regulation of PP synthesis includes the repressive EAR domain-containing Subgroup 4 MYB transcription factors^38^. All 4 members in Arabidopsis (MYB3, MYB4 MYB7 and MYB32) have been found to repress genes of the PP pathway. While MYB7 represses mainly flavonol biosynthesis^39^, MYB3, MYB4 and MYB32 target expression of *C4H*. We chose to employ MYB4, as this is the only member confirmed to bind directly to the promoter region of *C4H*^38^, to suppress PP synthesis specifically in the barrier tissues. To achieve this, we established plant lines containing MYB4 driven by promoters active in a transient peak in the differentiating endodermis (p*CASP1*)^40^ or in the suberizing endodermis and periderm (p*ELTP*)^7^ (Supplementary Figure 3d). Intriguingly, only plants with p*CASP1*-driven expression of MYB4 showed CS dysfunction (Figure 3a, d). This was manifested as a “discontinuous” CS with irregular patches (Figure 3a, b). In line with their characterized transcriptional targets, MYB3 and MYB32 showed similar effects, whereas no effects on CS formation was seen using MYB7 (Supplementary Figure 3a,c). No significant differences were found in any mutant root length when compared to WT (Supplementary Figure 3b). As this CS disruption could be complemented by addition of coniferyl alcohol to the media (Figure 3b, e), we conclude that the effect of MYB4 was specific to PP synthesis and not a more general interruption of endodermal differentiation, e.g. interference with MYB36 activity^41,42^. In support of this, plants with ectopic MYB4 expression in the differentiating endodermis showed unaltered localization of the CS protein CASP1^2^, fused to GFP, contrasting with the aberrant lignin depositions in the cell wall (Figure 3c). In summary, these experiments demonstrate that PP synthesis in the early endodermis is necessary for correct CS lignification and thereby function.

**Figure 3.**
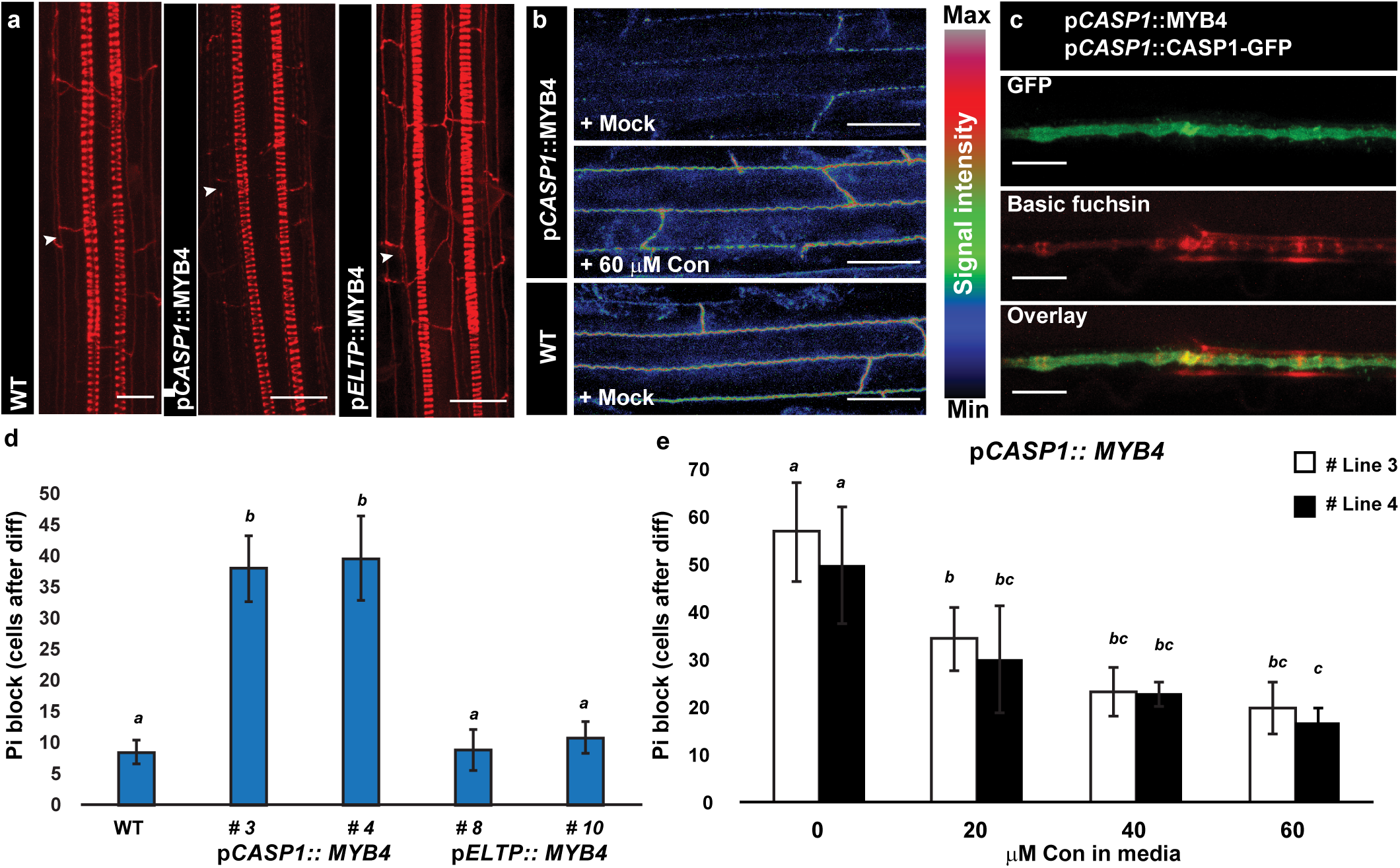
Endodermis specific expression of MYB4 alters lignin deposition and CS formation. a) Longitudinal maximum projections of lignin deposition in the endodermis and vasculature of 5-day-old seedlings. Lignin was highlighted using Basic Fuchsin staining stained using an established ClearSee-based protocol54. Arrowheads point toward the Casparian strips. The phenylpro-panoid-repressive transcription factor MYB4^38^ was expressed using promoters active in the differentiating endodermis (p*CASP1*)^40^ or mature endodermis and periderm (p*ELTP*)^7^. b) Surface view of endodermal CS stained by basic fuchsin in plants treated with either ethanol (mock) or coniferyl alcohol (Con) upon germination. c) Projected surface view of an endodermal cell with highlighted Casparian strip domain by CASP1-GFP and lignin deposition by Basic fuchsin staining in 5-day-old plants expressing MYB4 under the p*CASP1* promoter. d) Functional analysis of Casparian strip blockage by measurement of propidium iodide (PI) diffusion in the differentiating zone of the endodermis^37^ of individual lines with ectopic MYB4 expression in the endodermis. e) Blockage of PI in lines with p*CASP1*-driven MYB4 expression versus wildtype under increasing amount of coniferyl alcohol (Con). Plants were germinated and grown for 5 days in presence of the indicated substrate before measurement of PI penetration. N=6. All error bars are SD, letters refer to individual groups in a one-way ANOVA analysis with a post-hoc multiple group T-test (Tukey). Scale bars are 10 *μ*m.

The remaining CS-associated lignin observed in p*CASP1*::MYB4 plants could be caused by PP synthesis in adjacent tissues and/or compensatory mechanisms activated by the SGN-dependent surveillance system^15^. To address this, we created plants with p*CASP1*-driven MYB4 expression in combination with the *sgn3* mutant^13^. Interestingly, only in this mutant background could we observe a significant repression of *C4H* expression in whole roots (Figure 4a), which is likely due to the strong expression of *C4H* across endodermis-adjacent tissues (Figure 1). However, this might also indicate that MYB4 repression of *C4H* in the differentiating endodermis is partly alleviated through SGN-dependent activation^12^. In line with this, externally applied CIF2 peptide could only induce activity of the *C4H* transcriptional reporter in plants with an active SGN pathway, yet not in an endodermis specific manner (Figure 4 b,c). Thus, to ensure that these effects are endodermis-specific, we expressed a p*CASP1*::MYB4-GFP construct in a *cif1cif2* double mutant, which knock-out root ligands for the SGN pathway and phenocopies *sgn3*, but allow for activation of the SGN pathway by addition of CIF peptides to the media^11^. When grown under mock conditions, these lines showed a complete absence of basic fuchsin-stainable lignin in the CS, while CIF2 treatment led to non-connected endodermal lignin depositions in both the CS and in the endodermal cells of the suberized zone (Figure 4d, Supplementary Figure 3e). In both cases, xylem lignification remained unaffected (Supplementary Figure 3e). In conclusion, the endodermis autonomously produces lignin monomers required for CS formation without dependency on synthesis in the underlying xylem. This production is partially controlled by activation of *C4H* through the SGN pathway. Interestingly, all lines harboring p*CASP1*::MYB4 expression showed suberin deposition patterns similar to wildtype, albeit without CIF-dependent ectopic suberization in the young endodermis (Supplementary Figure 4b), which suggests that MYB4 can also repress suberin deposition.

**Figure 4.**
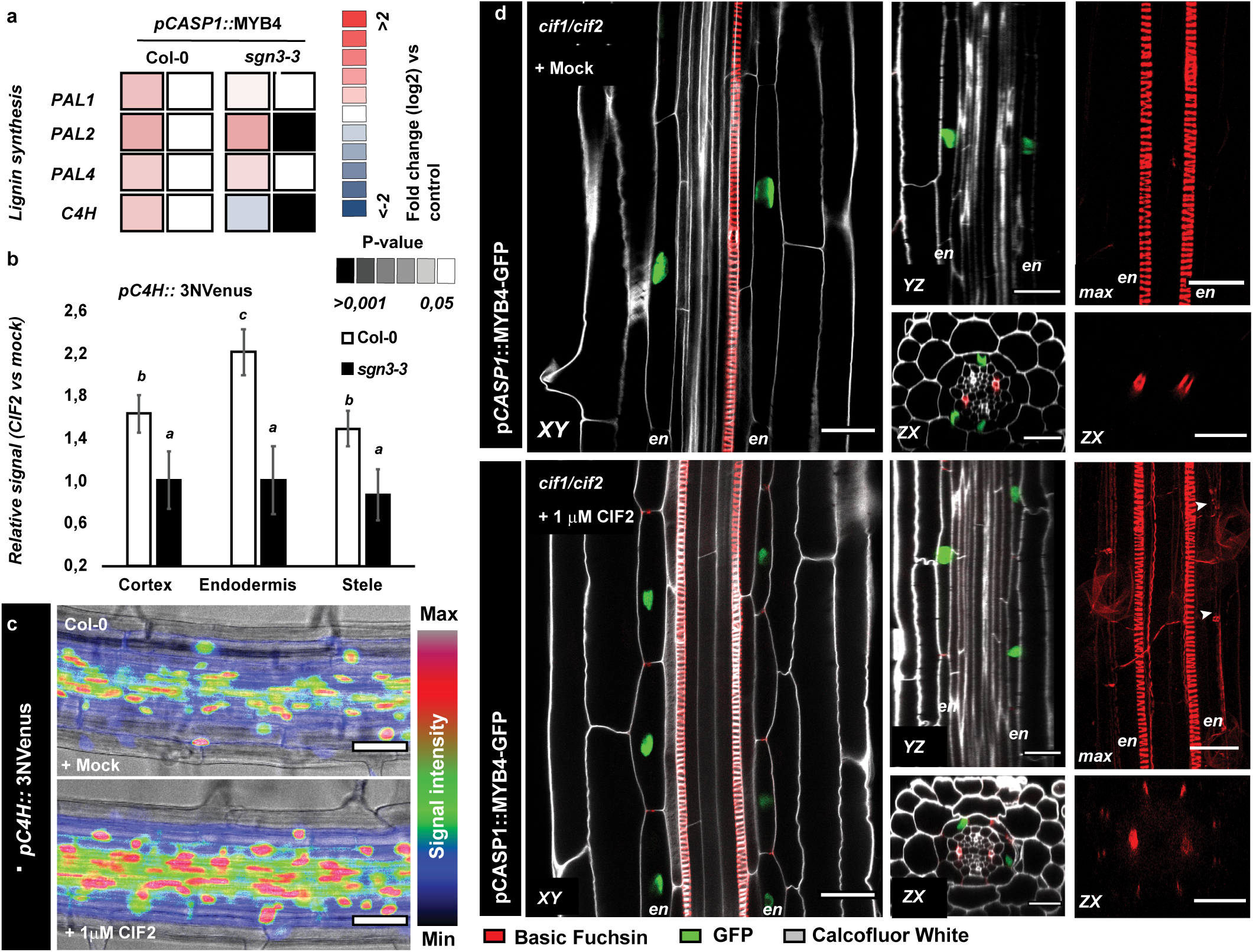
Compensatory SGN3-dependent lignification is activated upon MYB4 expression in the endodermis. a) Differential expression analysis of genes involved in the phenylpropanoid pathway in 5-day-old WT and *sgn3-3* plants expressing p*CASP1*::MYB4. Expression was normalized to each genotype without MYB4 expression. N= 3, P-values are based on a two-sided Students T-test (PA-vs mock-treated). b) Quantification of *C4H* activity in tissues of the differentiating endodermis using a transcriptional reporter (p*C4H*::3NVenus). Signals were normalized to expression of mock treated roots of the same line. All treatments were 24 h. N= 8. c) Representative image of *C4H* activity in the root differentiation zone of 5-day-old plants treated for 24 h with mock or 1 *μ*M CIF2 peptide d) Confocal projections of root sections expressing p*CASP1*-driven MYB4-GFP construct in the *cif1cif2* dko mutant upon treatment with mock or 1 *μ*M CIF2 peptide for 24 h. 6-day-old roots were fixed and stained with calcofluor white and Basic fuchsin^54^. Arrowheads point to non-connected lignification, en: endodermis. max; Maximum projection. All error bars are SD, scalebars are 10 *μ*m. Letters refer to individual groups in a one-way ANOVA analysis with a post-hoc multiple group T-test (Tukey).

### Continuous phenylpropanoid synthesis in the endodermis is essential for suberization

In line with MYB4 having a repressive effect on suberin, plants that continuously express MYB4 in the mature endodermis (p*ELTP* promoter) showed an almost complete loss of FY-stainable endodermal suberin (Figure 5a,b). MYB3 and MYB32 phenocopied this effect, whereas no decrease in suberin deposition was seen in p*ELTP*::MYB7 expressing plants (Supplementary Figure 3a). To confirm that this was due to a reduction in suberin content, we performed a GC-MS-based polyester analysis of p*ELTP*::MYB4 root fractions. Indeed, this revealed a strong decrease in all major aromatic and aliphatic suberin components when compared to wildtype levels (Figure 5e, Supplementary Figure 5a). To investigate if expression of MYB4 also affects pre-established suberin depositions, we used a p*ELTP* promotor-driven estradiol (E2D)-inducible^43^ system to express a MYB4-GFP fusion (p*ELTP*::XVE>>MYB4-GFP) (Figure 2c and f). Seedlings containing this construct were grown for 4 days to allow for suberin deposition on mock plates and then transferred to mock-or E2D-containing plates for an additional 2 days. Interestingly, after a total of 6 days, the suberized zone was strongly reduced in induced roots, indicating that a functional PP pathway is necessary to maintain a suberized barrier in the endodermis (Figure 5c). Neither coniferyl alcohol nor ferulic acid could revert this effect (Figure 5c), which can be explained by the fact that MYB4 represses other genes of the PP-pathway such as 4-Coumarate:CoA Ligases (4CLs), involved in ferulic acid-CoA conjugation^38^. To investigate this further, we performed a similar but reciprocal experiment with 4 days of germination on E2D-containing plates followed by 2 days recovery on mock-containing plates with EtOH or PP-metabolites. This recovery phase allowed MYB4-GFP degradation, as seen by disappearance of GFP signal in the mature endodermis (Figure 5d). Interestingly, in these experiments we observed an increase in FY-stained cells (Figure 5c), which was strongly increased specifically by addition of FA to the recovery plates (Figure 5c, Supplementary Figure 4a). This corroborates that FA and, importantly, not lignin monomers, are required for correct suberin deposition in the endodermis and is in line with our PA treatment analysis (Figure 2f). GC-MS analysis of endodermal suberin upon short-term or constant MYB4-GFP induction led to comparable reductions of both aromatic and aliphatic suberin components (Figure 5c and e, Supplementary Figure 5b). Moreover, the fusion of MYB4 and GFP had no effect on the repressive effect of MYB4 on suberin (Supplementary Figure 4c). Thus, the effect of our p*ELTP*::XVE>>MYB4-GFP construct on suberin recapitulates that of both PA treatment and the p*ELTP*::MYB4 lines, which strongly indicates that suberin is actively removed from the endodermis cell wall upon repression of PP metabolite production.

**Figure 5.**
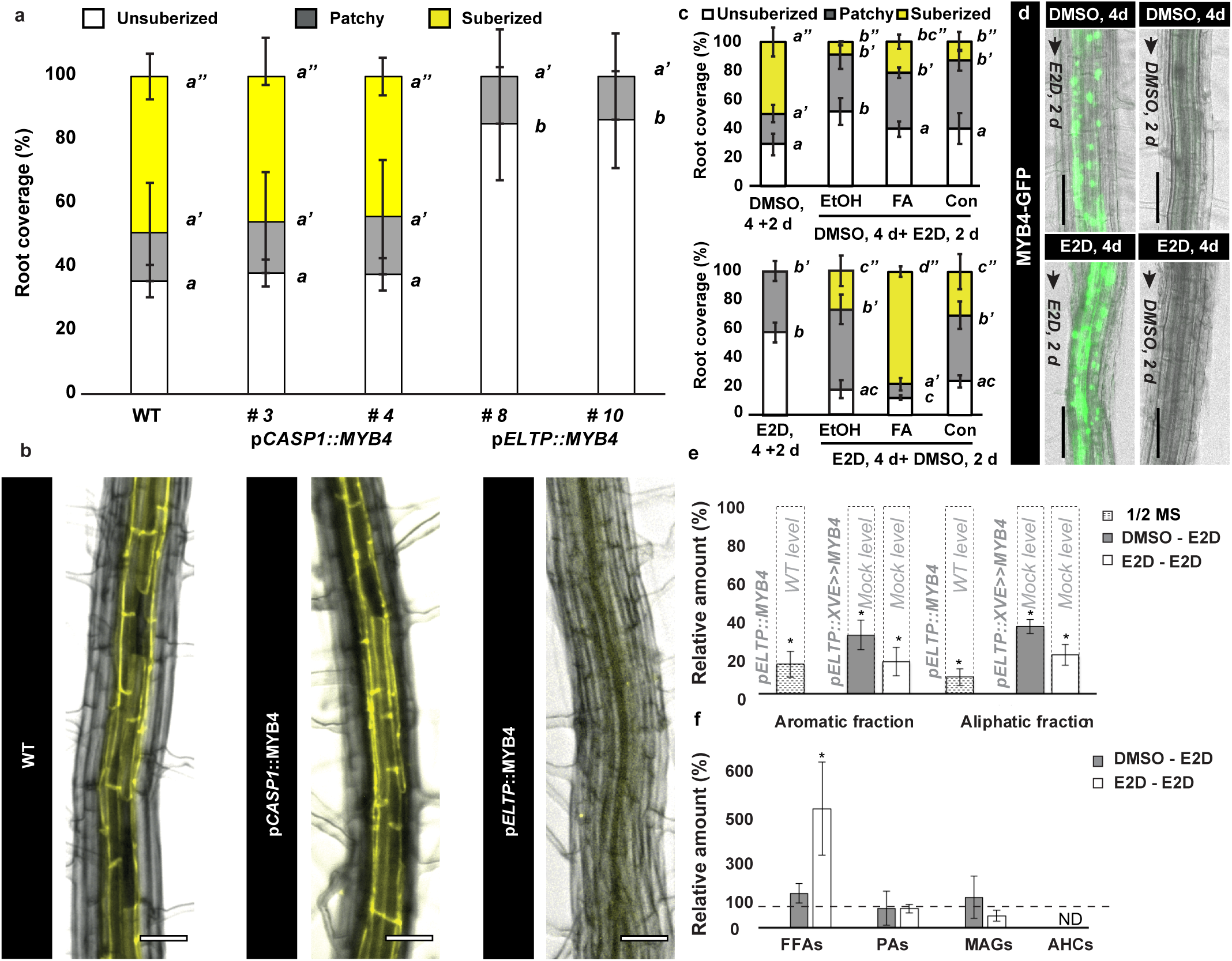
Repression of PP synthesis in the mature endodermis leads to suberin detachment. a) Quantification of Fluorol yellow (FY) staining in 5-day-old roots of lines expressing p*CASP1*-or p*ELTP*-driven MYB4. b) Representative FY-stained 5-day-old root from a) at similar developmental stages. Scale bars are 25 *μ*m. c) Plants expressing an beta-estradiol (E2D)-inducible MYB4 construct in the mature endodermis p*ELTP:*:XVE>> MYB4-GFP, were grown for 4 days on either DMSO-(upper) or 5 µM E2D (lower) -containing media and transferred to plates containing E2D or DMSO, respectively, in combination with ethanol (EtOH), 20 *μ*M Ferulic acid (FA) or 20 *μ*M Coniferyl alcohol (Con) for two days before suberin quantification by FY staining. d) GFP signal in 5-day-old p*ELTP*::XVE>>MYB4-GFP expressing plants grown under different regimes of DMSO or 5 *μ*M E2D. Scalebars are 50 *μ*m. e) Relative amount (%) of suberin aromatic and aliphatic fractions from 8-day-old p*ELTP*::XVE>>MYB4-GFP expressing roots under E2D treatment regimens (6 days mock + 2-days ED2: “DM-SO-E2D” or 6 days E2D + 2 days -E2D: “E2D-E2D”), WT and p*ELTP*::MYB4. Levels were normalized to WT or mock treated plants of identical age for p*ELTP*::MYB4 (white with stripes) and p*ELTP*::XVE>>MYB4-GFP (grey, white), respectively N= 3-5, * P< 0,05, T-test vs WT or mock. f) Relative amount (%) of chloroform extractives of the endodermis of 18-day-old p*ELTP*::XVE>>MYB4-GFP expressing roots under different E2D treatments (2-day, DMSO-E2D or 18-day, E2D-E2D) N= 3-4, * P< 0,05, T-test vs mock. AHCs: Alkyl hydroxycinnamates, FFAs: Free fatty acids, MAGs: Monoacylglycerol conjugates, ND: Not detected, PAs: Primary alcohols, WT: Wildtype. All error bars are SD. Letters refer to individual groups in a one-way ANOVA analysis with a post-hoc multiple group T-test (Tukey).

### Unbound aliphatic suberin constituents accumulate in absence of aromatics

The reduced levels of aliphatic components in the suberin of plants with p*ELTP*-driven MYB4 expression suggest that these components cannot be incorporated in the final suberin structure without aromatic components. Alternatively, this might be due to a negative feed-back loop in which absence of aromatics affect biosynthesis of aliphatic components. To clarify this, we tested whether free aliphatic components accumulate upon MYB4-GFP induction by performing a chloroform-based surface wax extraction^44^ of the endodermis-containing root parts. Surprisingly, only in continuously-induced plants did this reduction co-occur with a significant overaccumulation of aliphatic constituents, such as very-long-chain fatty acids, characteristic for suberin (P<0,01, Students T-test vs mock) (Figure 5f, Supplementary Figure 5c). This suggests that when no endodermal suberin deposition can be established due to limited PP metabolites, aliphatic constituents meant to be incorporated into the suberin matrix are likely accumulating as unpolymerised fatty acids in the apoplast. Upon short-term induction, these aliphatic constituents are either not soluble using our employed method, or they are degraded and utilized in other pathways.

### Repression of phenylpropanoid synthesis in periderm leads to collapse of the cork barrier

In 21-day-old roots undergoing secondary growth, we observed activity of all analyzed PP expression reporters in the cork and phellogen, at the “patchy” and mature stages of periderm formation (Figure 6a). The periderm might therefore also function tissue-autonomously to produce monomers for construction of the cork barrier. As the *ELTP* promoter is also active during secondary growth (Supplementary Figure 6a) we set out to address this question. Similar to the endodermis-specific analysis, 12-day-old roots of p*ELTP*::MYB4 expressing plants showed a reduction of lignin and suberin staining of cork cells when compared to wildtype (Supplementary Figure 6b-c). This effect was maintained through periderm development (Figure 6b and c and Supplementary Figure 6d). Moreover, these plants had collapsed cork cells (Figure 6c), which compromised the barrier function as visualized by increased toluidine-blue penetration (Figure 6c). Thus, the periderm appears to also be dependent on tissue-autonomous PP production in a similar manner to the endodermis. Next, we employed the p*ELTP*::XVE>>MYB4-GFP system to investigate if suberin in periderm and cork cell layers shows a similar dependency on continuous PP synthesis as the endodermis. To ensure onset of cork differentiation, we employed a matrix setup consisting of growth for 12 days on plates containing either mock or 5 *μ*M E2D with a consecutive transfer to mock or E2D for an additional 8 days to allow constant expression of MYB4-GFP, or induction in pre-formed peridermal cells. In both cases, we observed cork cell collapse similar to the p*ELTP*::MYB4 plants, however this appeared to be stronger in plants with constant induction (Figure 7a). After 8 days of E2d treatment, FY-stainable suberin was still detectable in the cork, albeit at lower levels than mock-treated plants (Figure 7a), whereas constant induction phenocopied the constitutive lines (Figure 7a). This likely reflects radial growth of the periderm, and suggest repression of *de novo* deposition in the developing periderm upon MYB4-GFP expression. To check this, and to analyze the fate of the suberin constituents in the periderm, we isolated the cork-containing root parts and subjected them to the same GC-MS-based pipeline as the endodermis samples. All cases (including constitutive p*ELTP*::MYB4 plants) had a strong reduction in suberin constituents when compared to the respective controls (Figure 7b and Supplementary Figure 6d and 7a). This is in line with ectopic MYB4 expression repressing *de novo* suberin deposition. When compared to mock or WT treatment, both constantly-, short-term-induced MYB4-GFP and constitutive MYB4 expression showed reduced content of alkyl hydroxycinnamates (AHCs) (Figure 7c and Supplemental Figure 7b-c). Although these aromatic-conjugated aliphatic compounds are typically found in low levels using this extraction method^44^ (Supplemental Figure 5c), this is likely due to the repressive effect of MYB4-GFP on aromatic constituents, which suggests that these originate from the cork. In contrast to the endodermis inductions, repression in the periderm led to increased accumulation of aliphatic suberin constituents in the chloroform fractions (Figure 7c, Supplementary Figure 7b-c). Surprisingly, constitutive repression using the XVE system led to a decrease in these components (Supplementary figure c), which is probably related to pleiotropic effects of the repressed periderm barrier formation. Based on these observations, we conclude that similar to the endodermis, suberin cannot be incorporated correctly into the cork barrier upon ectopic MYB4 expression. However, this also suggest that, upon short-term MYB4 induction, endodermal constituents might be re-directed into other pathways, while in cork, they over-accumulate as soluble constituents. Finally, to substantiate that ectopic MYB4-GFP expression in the cork and the consequential cell collapse give rise to a non-functional periderm barrier, we grew plants on mock-containing plates for 7 days followed by transfer to mock or 5 *μ*M E2D-containing plates with or without 100 mM NaCl for an additional 7 days. Only the combination of E2D (and the resulting MYB4-GFP expression) and NaCl led to a significant (P<0,01, pairwise T-test) increase in leaf chlorosis (Figure 7d and e and supplemental Figure 8a). This supports a role for the periderm in salt stress protection and emphasizes the defective barrier in our analysis. In all cases NaCl stress lead to reduced root lengths, without significant difference between the mock and E2D-treated plants (Figure 7e).

**Figure 6.**
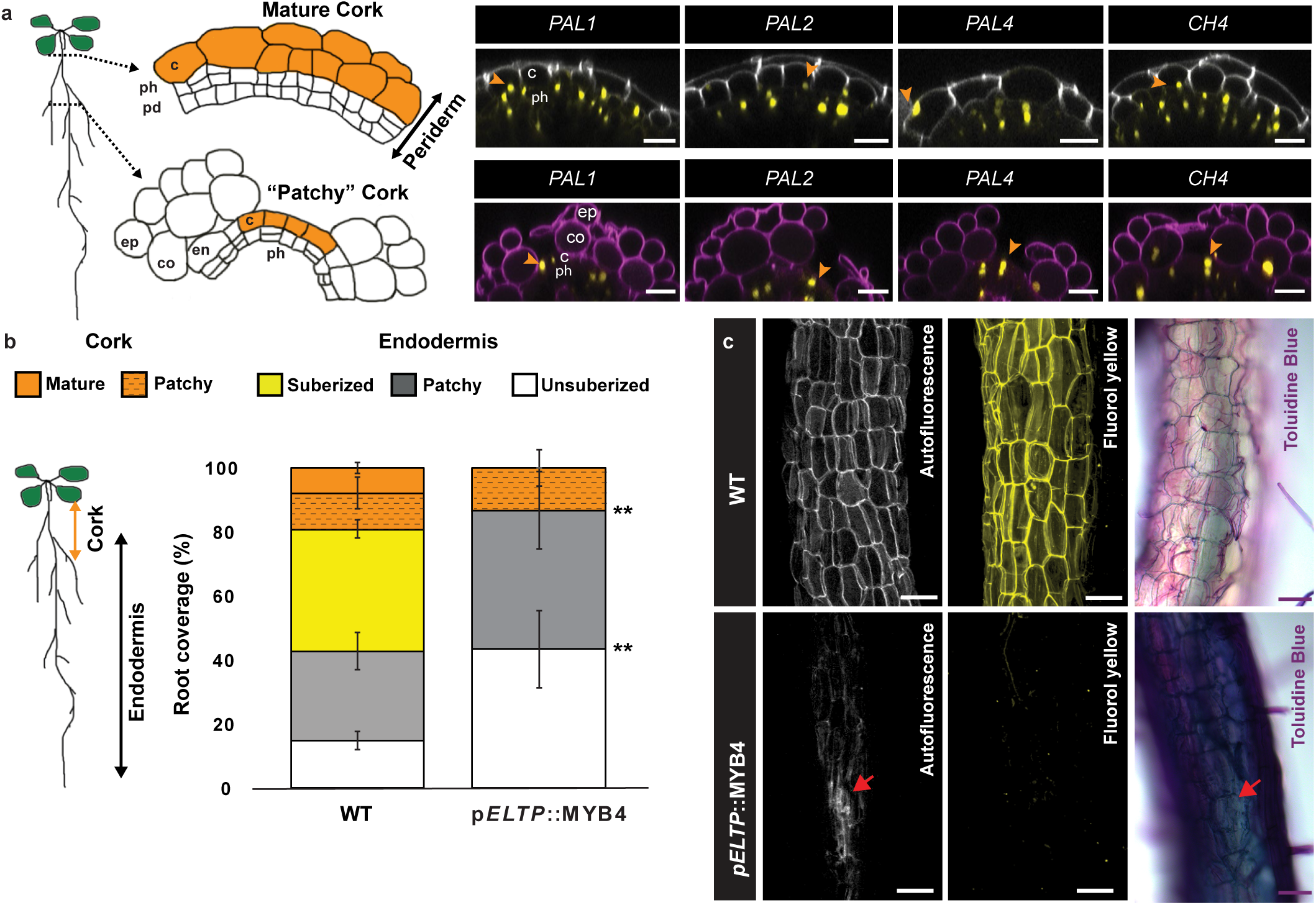
Collapse of cork formation by ectopic expression of MYB4 in the periderm. a) During secondary growth, the root increases in girth, the endodermis undergoes programmed cell death and its barrier function is eventually replaced by the periderm^16^. This is a developmental process, and zones of “patchy” and mature cork formation can be identified. Activity of a nuclear localized (NLS) 3x mVenus fusion reporters driven by promoter regions of root-expressed phenyl alanine lyase (*PAL*) and cinnamate 4-hydroxylase (*C4H*) homologues are shown in 21-day-old Arabidopsis roots at the different cork developmental stages. Cell walls were high-lighted using the intrinsic autofluorescence of cork cells (grey) or propidium iodide (magenta) for zones with “patchy” periderm areas. Orange arrowheads point cork cells with positive marker expression. Scalebars are 20 *μ*m. b) Quantification of suberin deposition in 12-day-old WT and p*ELTP*::MYB4 expressing roots using Fluorol Yellow (FY) staining 27 (N= 6 *; p< 0,05; ** p<0,01; Two-sided Students T-test vs WT). c) Autoflu-orescence (Left panel), FY staining (middle panel) and Toluidine blue (right panel) of cork in the periderm of wildtype (WT) and p*ELTP*::MYB4 expressing 19-d-old roots. Red arrows point to collapsed cork cells. Scalebars are 50 *μ*m. c: cork, co: cortex, ep: epidermis, pd: phelloderm, ph: phellogen WT: Wildtype. All error bars are SD

**Figure 7.**
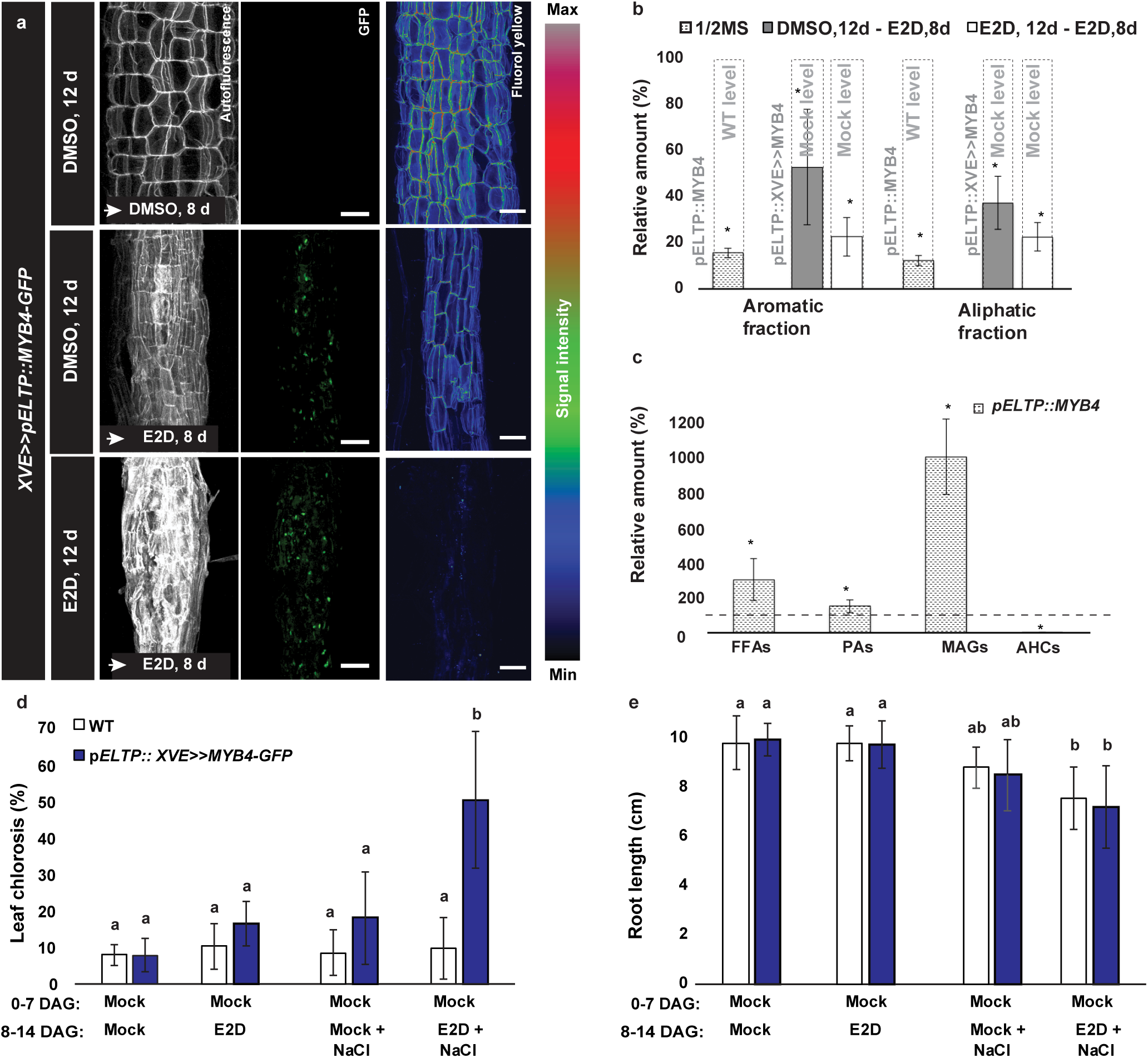
Cork barrier functionality requires an active PP pathway. a) Autofluorescence (left panel), GFP(middle panel) or Fluorol Yellow (FY) (right panel) signal in the cork of 20-day-old p*ELTP*::XVE>>MYB4-GFP roots under different mock and beta-estradiol (E2D) treatment regimes. b) Relative amount (%) of suberin aromatic and aliphatic fractions form the cork of 20-day-old WT, p*ELTP*::MY4 and p*ELTP*::XVE>> MYB4-GFP roots. Levels were normalized to WT and mock treated plants of identical age for p*ELTP*::MYB4 and p*ELTP*::XVE>>MYB4-GFP respectively. N=3-4 * P< 0,05, T-test vs WT or mock c) Relative amounts (%) of chloroform extractives of 20-day-old p*ELTP*:: MYB4 expressing roots. Normalization was to WT plants of identical age N= 3, * P< 0,05, T-test vs WT. d) Leaf chlorosis (%) of WT or p*ELTP*::XVE>>MYB4-GFP plants grown for 7 days (0-7DAG) under standard conditions (Mock) and transferred for 7 days to plates with combinations of mock, 5 µM E2D and 100 mM NaCl treatments for an additional 7 days (8-14DAG). e) Primary root length quanti-fication of plants from d). All error bars are SD. *: P< 0,05, **, P<0,01 in a Two-sided Students T-test vs WT. Letters refer to individual groups in a one-way ANOVA analysis with a post-hoc multiple group T-test (Tukey). Scalebars are 50 *μ*m. AHCs: Alkyl hydroxycinnamates, DAG: Days after germination, FFAs: Free fatty acids, MAGs: Mono-acyl glycerol conjugates, ND: Not detected, PAs: Primary alcohols, WT: Wildtype.

## DISCUSSION

In the Arabidopsis root, polymerization of lignin in the CS and xylem occurs rapidly and is initiated approximately 10-12 cells after onset of cell elongation^2,37^. In the xylem, polymerization of lignin can continue after cell death^45^, which indicates that monolignols can diffuse from surrounding cells. Most phenylpropanoids have physio-chemical properties that allow passive diffusion across membranes^46^, and it is appealing to consider if CS lignin originates from xylem, given the steles role in secreting barrier surveillance CIF peptides. Yet, a previous study reported (with low resolution) that *C4H* is expressed in the endodermis^47^, but did not demonstrate that it is involved in lignification of the CS. Our work clearly demonstrates that tissue-autonomous production of lignin monomers is required for CS formation. In line with this, the monolignol-specific ABCG29/PDR1 transporter is expressed in both the vasculature and endodermis^48^, which supports that active export of monolignols occurs in both cell-types. This compartmentalization might add additional spatial control to the PP synthesis and could thereby alleviate bottlenecks for production of individual compounds produced in either tissue from common substrates. Intriguingly, PP-metabolism aimed at producing monolignols and aromatic hydroxycinnamates appear to be restricted to certain developmental zones within the endodermis (Figure 3a and Figure 5a). In addition, we observed *PAL4* activity only in the suberizing zone of the endodermis (Figure 1), suggesting that this isoform might serve a role in production of distinct metabolites, possibly earmarked for suberin production. Future studies that include careful high-resolution expression pattern analysis of PP-pathway gene expression within the roots is therefore likely to reveal currently unappreciated regulatory mechanisms that drive cell-type and developmental stage specific PP-metabolic pathways within the root.

Our work further show that not only are PP-derived metabolites components of the suberin matrix, they serve to stabilize and anchor the polymers to the cell wall and regulate active turn-over of suberin in the endodermis. Reduced availability of Fe, Zn or P leads to decreased endodermal suberin deposition, particularly, in the xylem pole^27^. An active mechanism for turn-over of suberin would therefore allow the plant to quickly respond to abiotic changes in the external environment. Moreover, symbiotic associations with for example arbuscular mycorrhiza-forming fungi exchange photosynthates from the plant in the form of sugars and fatty acids^49^ for assimilated phosphate, which could be correlated with suberin dynamics. The GDSL lipase Cuticle destruction factor 1 (CDEF1) involved in degradation of cutin^50^, has been employed as a tool to remove suberin in the endodermis^7,13,27,37^ illustrating the capacity of plants for suberin catabolism by a rather simple mechanism. Our setup with inducible repression of PP synthesis and the corresponding turn-over of suberin, provides an excellent platform to unravel such putative aspects of suberin function in inter-species associations.

Short-term repression of PP in the periderm led to release of multiple different classes of aliphatic compounds from the cork, (Figure 7c). Similarly, in the endodermis, only in constitutive MYB4 expression conditions did we see specifically a release of FAs. This is not due to a different relative suberin composition of these tissues (Supplementary Figure 8b), but supports that suberin dynamics is different in cork and endodermal tissues. Our analysis is surface oriented, thus extraction from the periderm, an outer tissue, is likely more efficient than from the endodermis, as it is surrounded by other tissue layers. Still, waxes in the form of FAs can be released upon PP repression. This is in line with a previous observation that in young *in vitro* cultured roots most FAs are not incorporated into the suberin polymer, but exists as associated waxes^44^. These compounds might therefore rather reflect the pool of soluble aliphatic constituents available due to a continuous turn-over of suberin in the endodermis.

In conclusion, our work illustrates that genetic repressors can be used as tools to elucidate PP tissue-specific functions, providing a novel approach to affect root barrier formation with an unprecedented resolution. We demonstrate that the endodermis is self-sufficient in producing metabolites for establishment of the CS as well as a coherent suberin barrier. Our data support that continuous production of hydroxycinnamates such as ferulic acid is essential for suberin attachment in the endo-and peridermal cell walls. We thus provide a novel genetic tool that can be employed to tease apart and gain insight into the independent functions of PP synthesis at different timepoints and in different cell layers, circumventing pleiotropic effects on xylem formation.

## MATERIALS AND METHODS

### Plant material and growth conditions

For all experiments, Arabidopsis thaliana (ecotype Columbia 0) was used. For p*CASP1* and p*ELTP* lines driving MYB4, the background contained a previously described p*GPAT5*:: mCitrin-SYP122 reporter. The T-DNA tagged lines *sgn3-3* (SALK_043282) and *cif1cif2* were previously described^11,13^. The p*CASP1*::MYB4 construct was independently transformed into each mutant background using the floral dip method. For observations and histological analysis, seeds were kept for 2 days at 4°C in the dark for stratification, then grown for 5 days at 22°C under 16 h light/8 h dark vertically on solid half-strength Murashige–Skoog (MS) medium without sucrose. For all periderm and GC-MS analysis, plants were grown under continuous light conditions.

### Plasmid construction

To generate endodermis specific expression constructs the genomic region of MYB3, 4 7 or 32 were amplified using specific primers (Supplementary table 1) and Gateway cloned into a pDONR221 entry vector using BP clonase II (Invitrogen) according to manufactures description. In case of MYB4-GFP, the genomic region was directly fused to GFP by overlapping PCR and inserted as one into the pDONR 221 vector. Together with previously generated P4L1r pDONR entry vectors containing either p*CASP1*, p*ELTP* or p*ELTP:*:XVE>> 27, the MYB entry vectors were recombined using LR-clonase II (Invitrogen) into a fastred selection-contain destination vector. For promoter constructs p*PAL1* (4038bp) p*PAL2* (3456bp), p*PAL4* (2332 bp) and p*C4H* (2450bp) were cloned into a modified gateway vector containing an attL4 R1r cassette cut with XbaI using Infusion cloning (Takara). The expression constructs were transformed into Col-0 or mutant backgrounds using the floral dip method^51^ and selected using FastRed selection^52^.

### GC-MS

For periderm analysis, sections of ∼2 cm of periderm from 20-day-old plants were collected directly from plates. Most lateral roots were cut off from the samples. For endodermis analysis, the whole root was collected if no periderm was already formed, or if the periderm was already present, only the lower portion of the root was collected. In both cases, the samples were washed by submerging three times in deionized water and carefully dried on paper towel before subsequent analysis. For suberin analysis, we followed a previously established protocol^21^. Briefly, the roots were treated with an enzymatic solution (Cellulase, Pectinase-Sigma-Aldrich, Germany) for 7 days. The solution was exchanged four times. After enzymatic treatment, unbound lipids were extracted from the roots by Soxhlet extraction with chloroform: methanol (1:1; v/v) for 2 days. The samples were dried and weighed. Consecutively, suberin was depolymerized using a 10% BF3/MeOH-based procedure^5^. In order to quantify (µg/mg sample) the suberin monomers detected by gas chromatography-coupled mass spectrometry (GC-MS), after the depolymerization, 25 µl of C32 alkane internal standard (13,5 mg/50 ml) was added in each sample. After the extraction, samples were concentrated by evaporation with N_2_ until ∼50 µl and posteriorly derivatized with 20 μL of BSTFA (bis-(N,O-trimethylsilyl)-tri-fluoroacetamide, Macherey-Nagel, Germany) and 20 μL Pyridin for 40 min at 70°C.

For chloroform-extraction analysis, fresh samples were used. After washing with deionized-water samples were carefully weighed. In order to quantify chloroform extracts (µg/g fresh weight), 22 μL of C15 (20 µl /50 ml) was added to each sample. following this, samples were submerged in chloroform for 90 sec^44^. The chloroform extracts were then evaporated with N_2_ until ∼50 µl. Finally, before GC-MS analysis, samples were derivatized with 20 μL of BSTFA (bis-(N,O-trimethylsilyl)-tri-fluoroacetamide, Macherey-Nagel, Germany) and 20 μL Pyridin for 40 min at 70°C. All extracts were measured using a Shimadzu TQ8040 GC-MS setup using splitless injection mode on a SH-Rxi-5SIL-MS column (30 m, 0.25 mm internal diameter, 0.25 µm film, Shimadzu Cooperation). The starting temperature was 50°C for two minutes, with an increase of 10°C per minute until 150°C, 150°C for one minute, with an increase of 3°C per minute until 310°C, and 310°C for 15 minutes. Helium was used as carrier gas with a flow rate of 0.86 ml min-1. The mass spectrometer was operated in electron impact ionization (EI) scan mode. The analyses were conducted with three or five replicates. Unless otherwise stated, statistical analyses were performed with IBM SPSS Statistics version 25 (IBM). First, the datasets were for homogeneity of variances using the Levene’s Test. The significant differences between two datasets were calculated using a Welch’s t-test in case of a non-homogenous variance or a Student’s t-test if the variance was homogeneous. For multiple sample comparisons, a one-way ANOVA with Tamhane’s post hoc (equal variance not assumed) or a Bonferroni correction (equal variance assumed) was performed unless otherwise stated. In the chloroform extractive analyses the blank was subtracted from C16 and C18 acids (all the other compounds were not detected in the blank). As C16 acid after blank subtraction was below detection it was not included in the analyses.

### Confocal microscopy

Confocal pictures were obtained using Leica SP8 or Zeiss LSM 880 confocal microscopes.

The excitation and detection window settings to obtain signal as follows: When using Leica SP8 (excitation, detection window), GFP (488 nm, 500–550 nm), mVenus, (514 nm, 518–560 nm), when using Zeiss LSM880 (excitation, detection window), Calcofluor White (405 nm,425-475 nm). All periderm pictures were acquired with a Zeiss LSM880 with the following settings: cork autofluorescence (405nm; 420-460nm), PI and Basic Fuchsin. (566 nm; 570-630 nm). FY (488 nm, 490-540 nm), GFP (488 nm,490-510 nm), Venus: (514 nm;. 520–540 nm). 3D reconstructions and Orthogonal views of a Z-stack were obtained using the ZEN Black software.

### Stainings

ClearSee staining coupled to cell wall staining were performed as recently described in ^53,54^ Briefly, plants were fixed in 3 mL 1 x PBS containing 4% p-formaldehyde for 1 hour at room temperature in 12-well plates and washed twice with 3 mL 1 x PBS. Following fixation, the seedlings were cleared in 3 mL ClearSee solution under gentle shaking. After overnight clearing, the solution was exchanged to new ClearSee solution containing 0.2% Fuchsin and 0.1% Calcofluor White for lignin and cell wall staining respectively. The dye solution was removed after overnight staining and rinsed once with fresh ClearSee solution. The samples were washed in new ClearSee solution for 30 min with gentle shaking and washed again in another fresh ClearSee solution for at least one overnight before observation. For quantification of Basic Fuchsin signal, recovering roots were normalized to signals from identically stained, non-PA treated Col-0 plants and plants taken directly from PA-containing plates (timepoint 0). For Fluorol Yellow staining of Arabidopsis root suberin vertically grown 5-day old seedlings were incubated solution of Fluorol Yellow 088 (0.01%, in lactic acid) and incubated for 30 min at 70 degrees. The stained seedlings were rinsed shortly in water and transferred to a freshly prepared solution of aniline blue (0.5%, in water) for counterstaining. Following this, seedlings were washed for 2-3 min in water and transferred to a chambered cover glass (Thermo Scientific), and imaged either as described above or using a Leica DM5500 wide field microscope (GFP filtercube ex: 470 nm/40 em:525/50 bs: 500). For CLSM fluorescence, Fluorol Yellow was detected using 488 nm as excitation wavelength, and collection of emission from 500-550nm. PI assays were done as described2 shortly, seedlings were incubated in water containing 10 μg /mL PI for 10 min and transferred into fresh water. The number of endodermal cells were scored using a Leica DM5500 wide field microscope (TX2 filtercube ex: 560 nm/40 em:645/75 bs: 595) from onset of cell elongation (defined as endodermal cell length being more than two times than width in the median, longitudinal section) until PI could not penetrate into the stele. Toluidine Blue penetration assay in the periderm assay was performed as described for analysis in the lateral root cap^55^. Briefly, plants were collected in water in a 6-well plate, transfer in a 6-well plate containing 0.05% Toluidine Blue, incubate for 2 min and washed twice with water prior mounting then them in water on a slide. Imaging was performed straight after mounting with a Zeiss Axiophot microscope. Chlorosis of leaves was measured in ImageJ56 by measuring green channel values images as previously described ^57^.

### qPCR analysis

Seedlings were grown on half MS without sucrose for 5 days. Only root parts (around 100 mg) were collected from each genotype and total RNA was extracted using a Trizol-adapted ReliaPrep RNA Tissue Miniprep Kit (Promega). Reverse transcription was carried out with PrimeScript RT Master Mix (Takara). All steps were done in at least triplicates and as indicated in manufactures’ protocols. The qPCR reaction was performed on an Applied Biosystems QuantStudio3 thermocycler using a MESA BLUE SYBR Green kit (Eurogentech). All transcripts are normalized to Clathrin adaptor complexes medium subunit family protein (AT4G24550) expression. All primer sets are indicated in Supplementary table 1

## ACKNOWLEDGEMENTS

We wish to thank Magdalena Marek and Colleen Drapek for their readthrough and insightful comments to the manuscript. We thank Dagmar Ripper for helping in the cork staining, and Robertas Ursache and Joop Vermeer for insightful discussions. The authors are grateful to the central imaging facility (CIF) at university of Lausanne for aid in image quantification and analysis. Work in the Ragni lab is supported by the Deutsche Forschungsgemeinschaft (DFG grant RA-2590/1-2).

## AUTHOR CONTRIBUTIONS

TGA acquired and interpreted data, conceived the research and wrote the manuscript. DM conducted and analyzed GC-MS data together with JK. RF interpreted data and contributed to writing the manuscript. DM and LR acquired the cork data. LR conceived the research, acquired and interpreted data and contributed to writing the manuscript. NG conceived the research and contributed to writing the manuscript. All authors have approved the final manuscript and agree to be personally accountable for the individual contributions.

## COMPETING INTERESTS

The authors declare no competing interest.

---

**Supplementary Figure 1.**
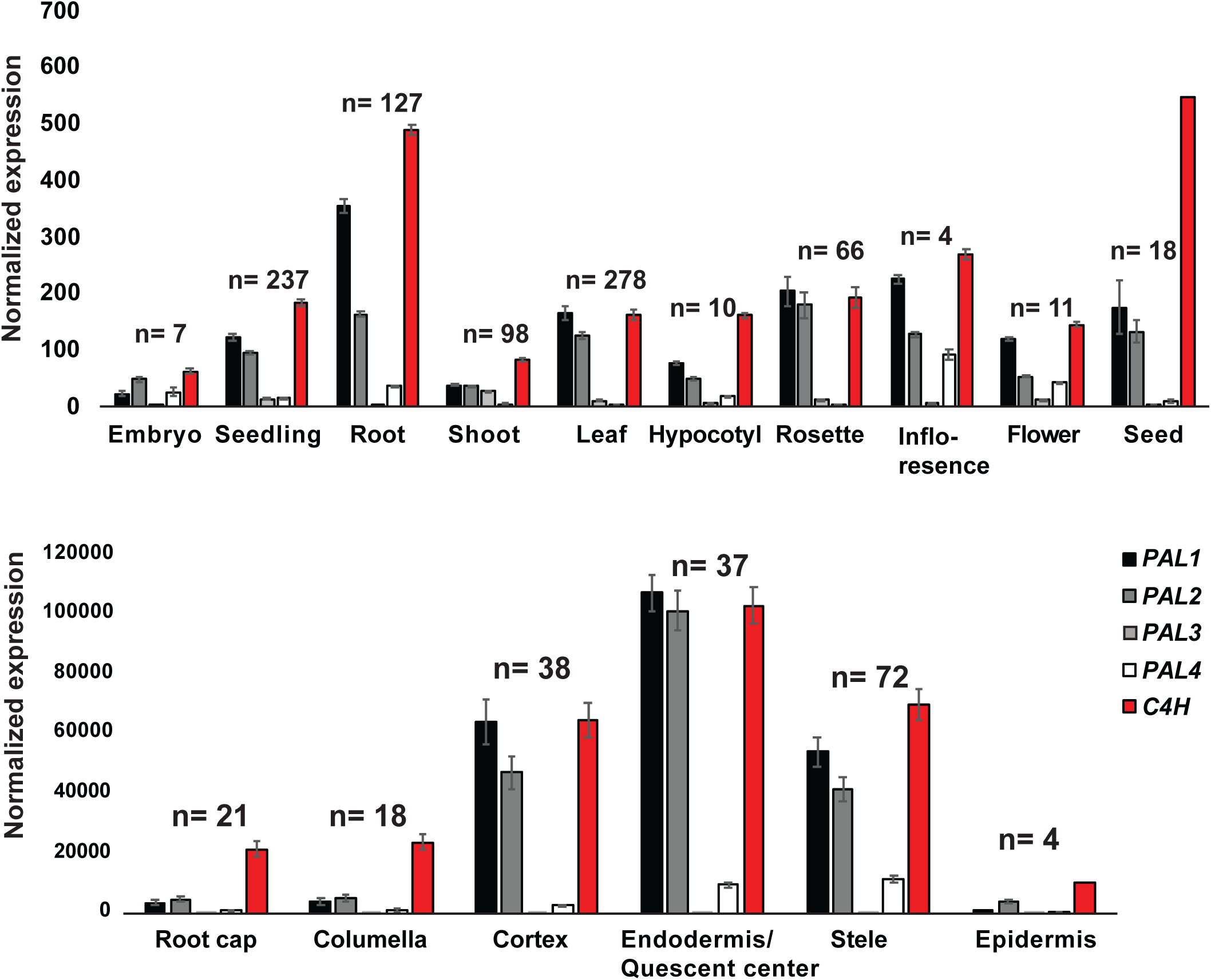
In silico expression analysis of essential phenylpropanoid synthesis genes. Arabidopsis tissue expression data based on publicly available dataset (Genevestigator)^35^ of genes involved in the phenylpropanoid pathway. The upper graph depicts different tissues while the lower shows expression across the root cell layers

**Supplementary Figure 2.**
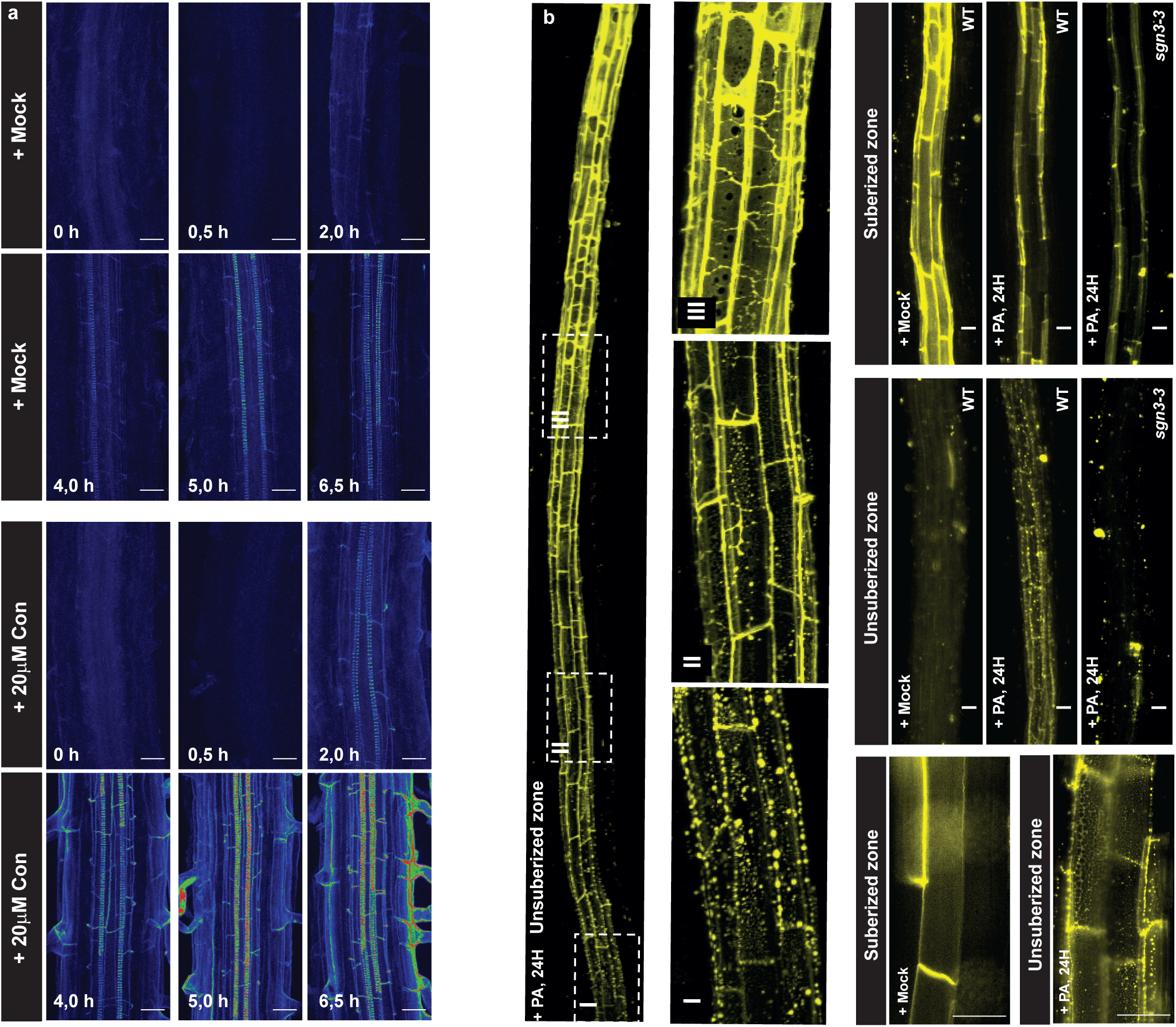
Behavior of lignin and suberin in roots upon PA treatment. a) Time course analysis of Basic fuchsin signal recovery in the non-lignified zone of xylem or Casparian Strip Domain (CSD) after 24 h of piperonic acid (PA) pre-treatment. 6-day-old plants were moved from PA-containing plates to recovery plates with either a mock solution or 20 *μ*M coniferyl alcohol (Con) at the indicated timepoints. Lignin deposition highlighted by Basic fuchsin signal in the endodermis and vasculature using an established ClearSee-based protocol^54^. b) Deposition of suberin visualized by Fluorol yellow (FY) in root zones of 5-day-old plants treated with mock or 1 *μ*M PA for 24 hours.

**Supplementary Figure 3.**
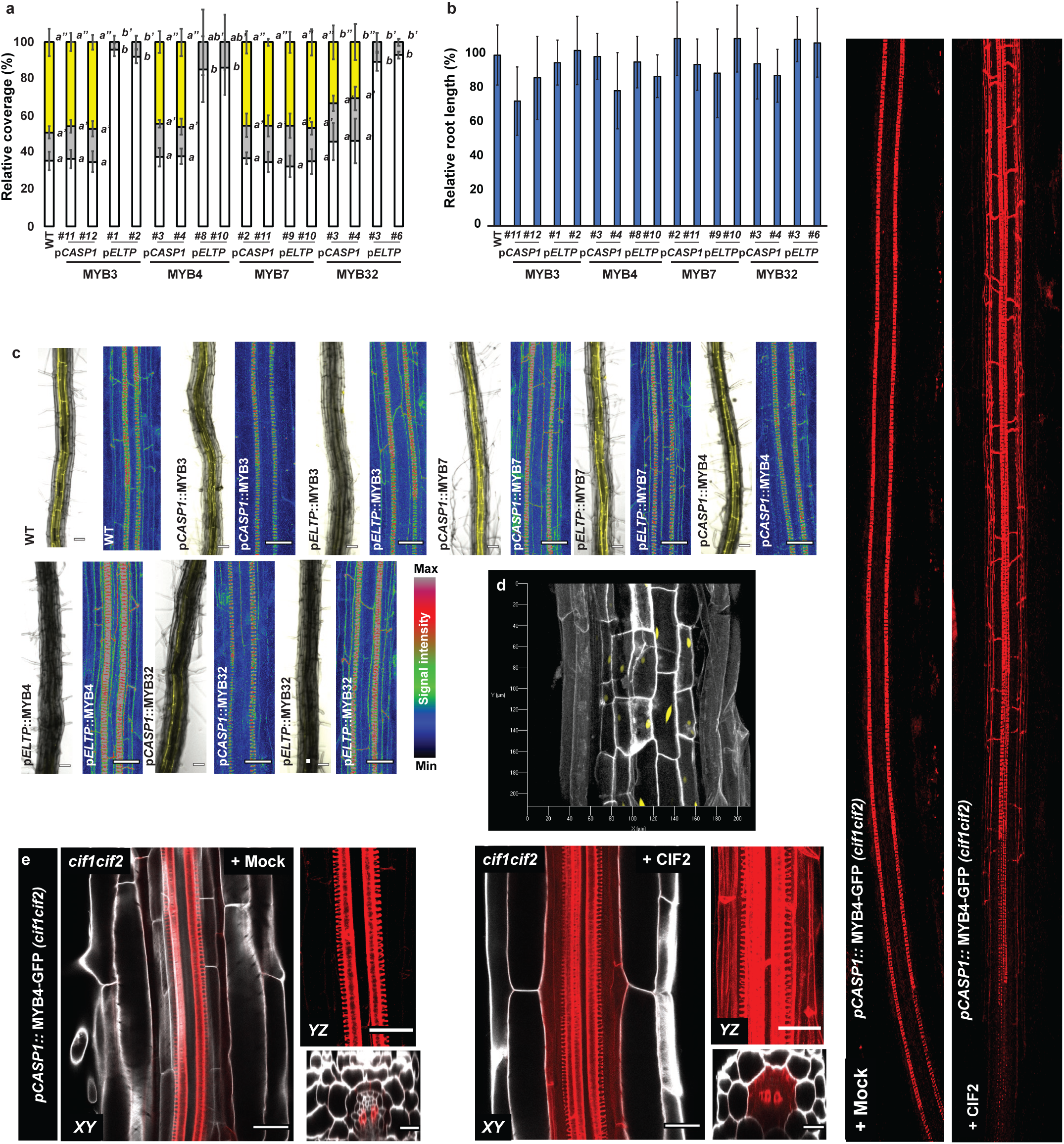
Lignin and suberin analysis upon ectopic expression of MYB-repressors. Quantification of Fluorol yellow (FY) positive suberin zones in 5-day-old roots of lines with p*CASP1*-or *pELTP*-driven MYB4 expression n=6. a) Root length relative to Wildtype plants subject to FY staining c) Suberin and lignin deposition highlighted by FY and basic fuchsin respectively, in the endodermis and vasculature of 5-day-old seedlings. The phenylpropanoid-repressive transcription factors were expressed using promoters active in the differentiating (p*CASP1*) or mature endodermis and periderm (*pELTP*). Scale bars are 25 *μ*m. d) Expression of pELTP:: NLS 3xm-Venus in the periderm of 20-day-old roots stained with Calcofluor white e) Confocal projections of root sections in the suberizing zone harvesting expression of a pCASP1-driven MYB4-GFP construct in the cif1cif2 dko mutant upon treatment with mock or 1 *μ*M CIF2 peptide for 24 h. 5-day-old roots were fixed and stained with calcofluor white and Basic Fuchsin using a previously established ClearSee-based prototol^54^. Scalebars are 10 *μ*m. All error bars are SD. Letters refer to individual groups in a one-way ANOVA analysis with a post-hoc multiple group T-test (Tukey).

**Supplementary Figure 4.**
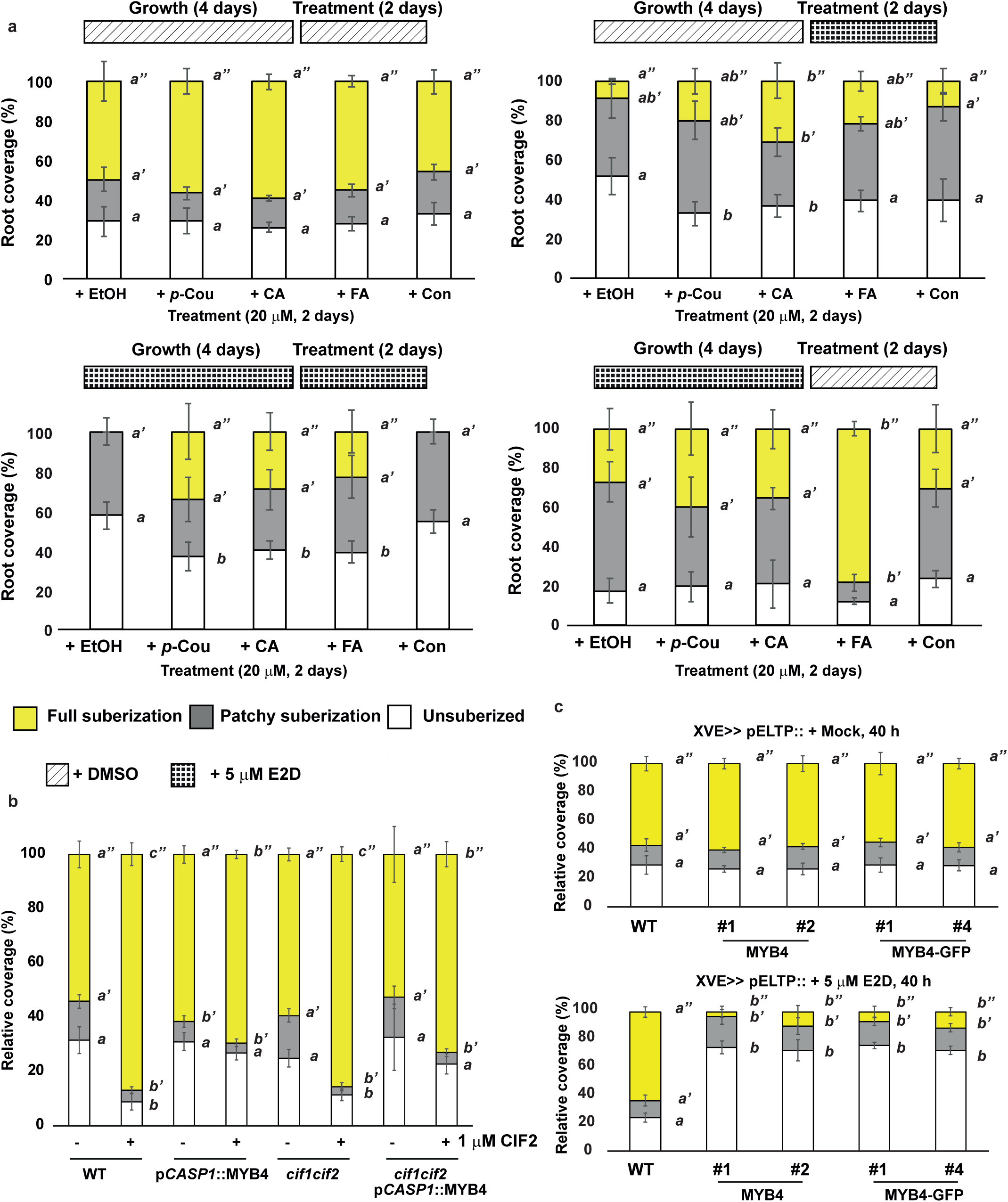
Fluorol yellow analysis of mutants and treatments. a) Quantification of fluorol yellow (FY)-stained endodermal suberization in 5-day-old Arabidopsis pELTP::XVE>>MYB4-GFP seedlings treated with either DMSO or 5 *μ*M beta-estradiol (E2D) in combination-regimes with mock (EtOH), 20 *μ*M p-coumaric acid (p-Cou) 20*μ*M caffeic acid (CA), 20 *μ*M ferulic acid (FA), or 20 *μ*M coniferyl alcohol (Con) for 48 h. b) Quantification of FY-stained endodermal suberization in 5-day-old WT, or *cif1cif2* dKO Arabidopsis plants with or without p*CASP1*::MYB4 expression. Plants were treated for 24 h with wither mock (H2O) or 1 *μ*M CIF2 peptide. c) Quantification of FY stained endodermal suberization in 5-day-old Arabidopsis p*ELTP*::XVE>>MYB4-GFP or MYB4 seedlings treated with either DMSO or 5 *μ*M E2D. For all experiments, N=6. All error bars are SD, Letters refer to individual groups in a one-way ANOVA analysis with a post-hoc multiple group T-test (Tukey).

**Supplementary Figure 5.**
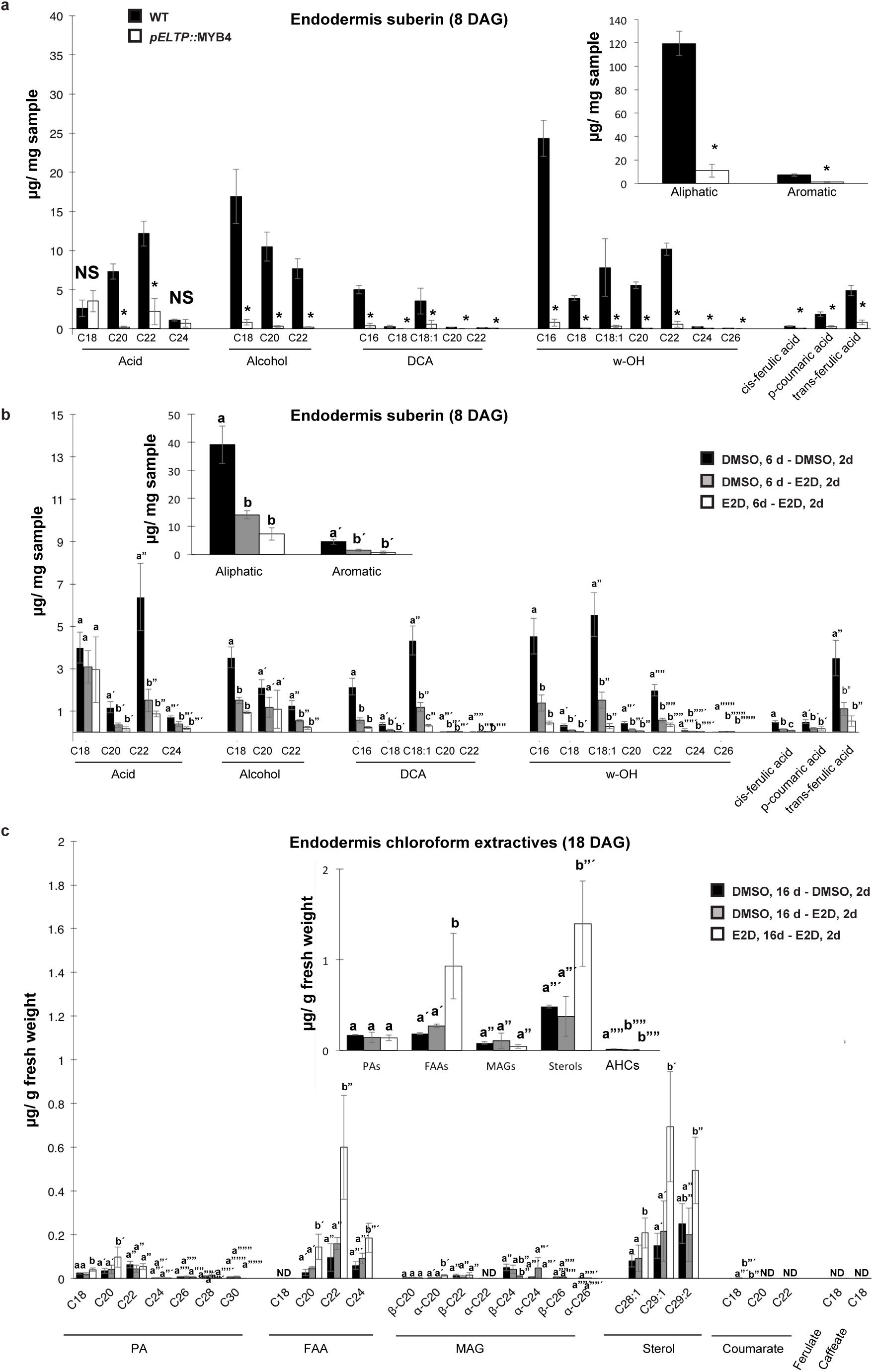
Suberin and Chloroform extractives quantification in the endodermis. a) Suberin quantification by Gas chromatography-coupled mass spectroscopy (GC-MS)-based analysis from the endodermis of 8-days-old wildtype (WT) and p*ELTP*::MYB4 expressing roots (N=4-5; T-Test *: P<0.05. b) Suberin quantification by GC-MS-based analysis from the endodermis of 8-days-old p*ELTP:*:XVE>>MYB4-GFP expressing roots under different beta-estradiol (E2D) treatments (One-way ANOVA (CI95%, N=3-4) c) Chloroform extractives quantification by GC-MS-based analysis from the endodermis of 18-day-old p*ELTP*::MYB4 expressing roots under different E2D treatment regimens (One-way ANOVA (CI95%, N=3). For the comparison of the suberin fractions, each compound was normalized to the total aliphatic or aromatic respectively. Numbers refer to carbon chain lengths of aliphatic fatty acid derivatives. DCA: *α–ω*-dicarboxylic acid, ND: Not detected, NS: Not significant. All error bars are SD.

**Supplementary Figure 6.**
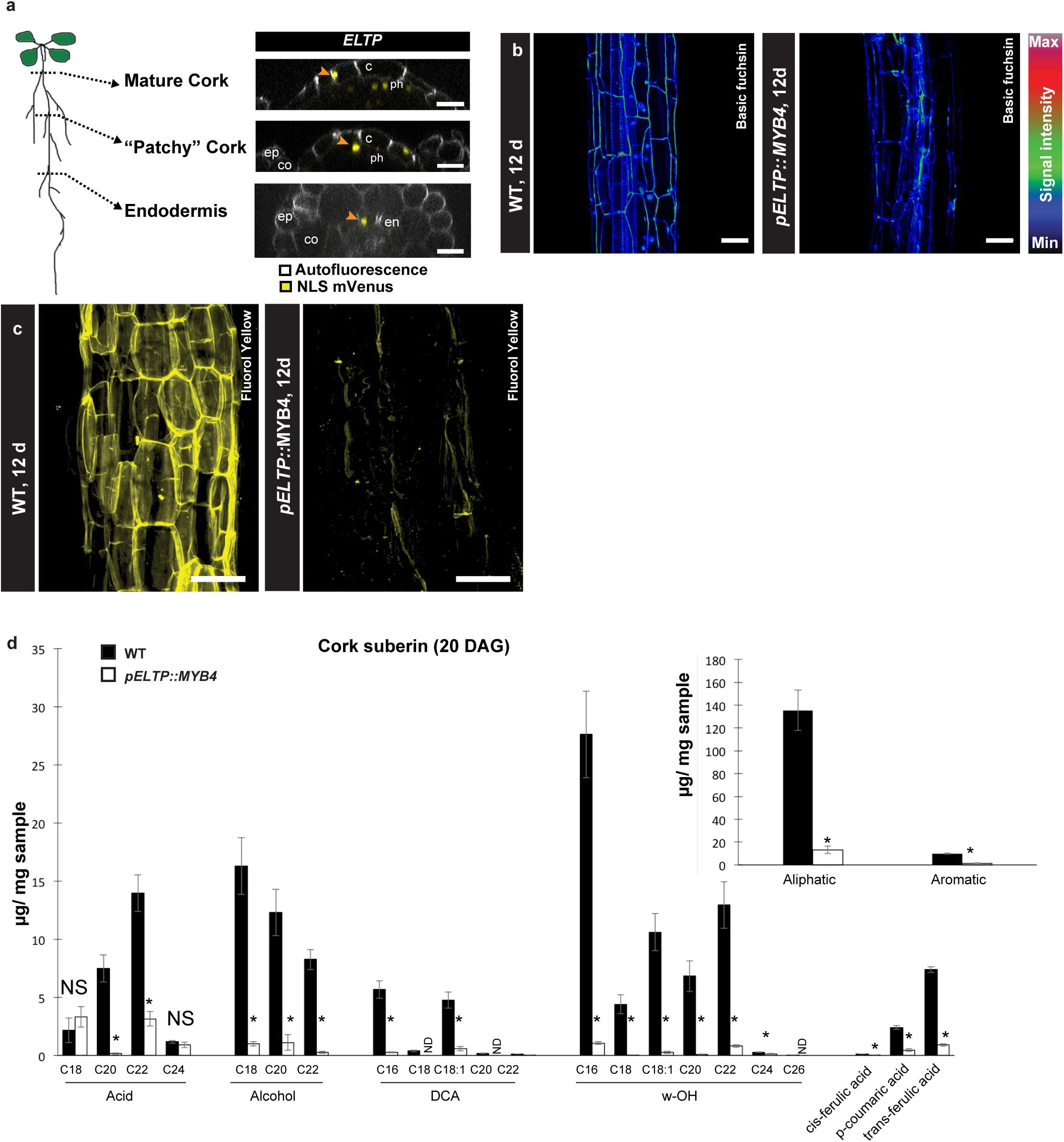
Cork phenotypes of pELTP::MYB4 roots. a) Representative images of p*ELTP*-driven nuclear-localized (NLS) 3x mVenus reporter in the mature and “patchy” periderm as well as endodermis of 14-day-old roots. Orange arrowheads highlight cells with p*ELTP* activity. Grey represent autofluorescence. Scalebars are 20 *μ*m. b) 12-day-old roots of WT and pELTP:MYB4 expressing plants were fixed and stained Basic fuchsin using a previously established ClearSee-based protocol. Scalebars are 20 *μ*m. c) Fluorol yellow (FY) staining of suberin in the cork of 12-day-old roots of WT and p*ELTP*:MYB4 expressing plants. Scalebars are 50 *μ*m. d) Suberin quantification by Gas chromatography-coupled mass spectroscopy (GC-MS)-based analysis from the cork of 20-day-old wildtype (WT) and p*ELTP*::MYB4 expressing roots (N=3-4; T-Test *: P<0.05). For the comparison of the suberin fractions, each compound was normalized to the total aliphatic or aromatic respectively. Numbers refer to carbon chain lengths of aliphatic fatty acid derivatives., c: cork, co: cortex DCA: *α–ω*-dicarboxylic acid, ep: epidermis, NS: Not significant, ph: phellogen. All error bars are SD.

**Supplementary Figure 7.**
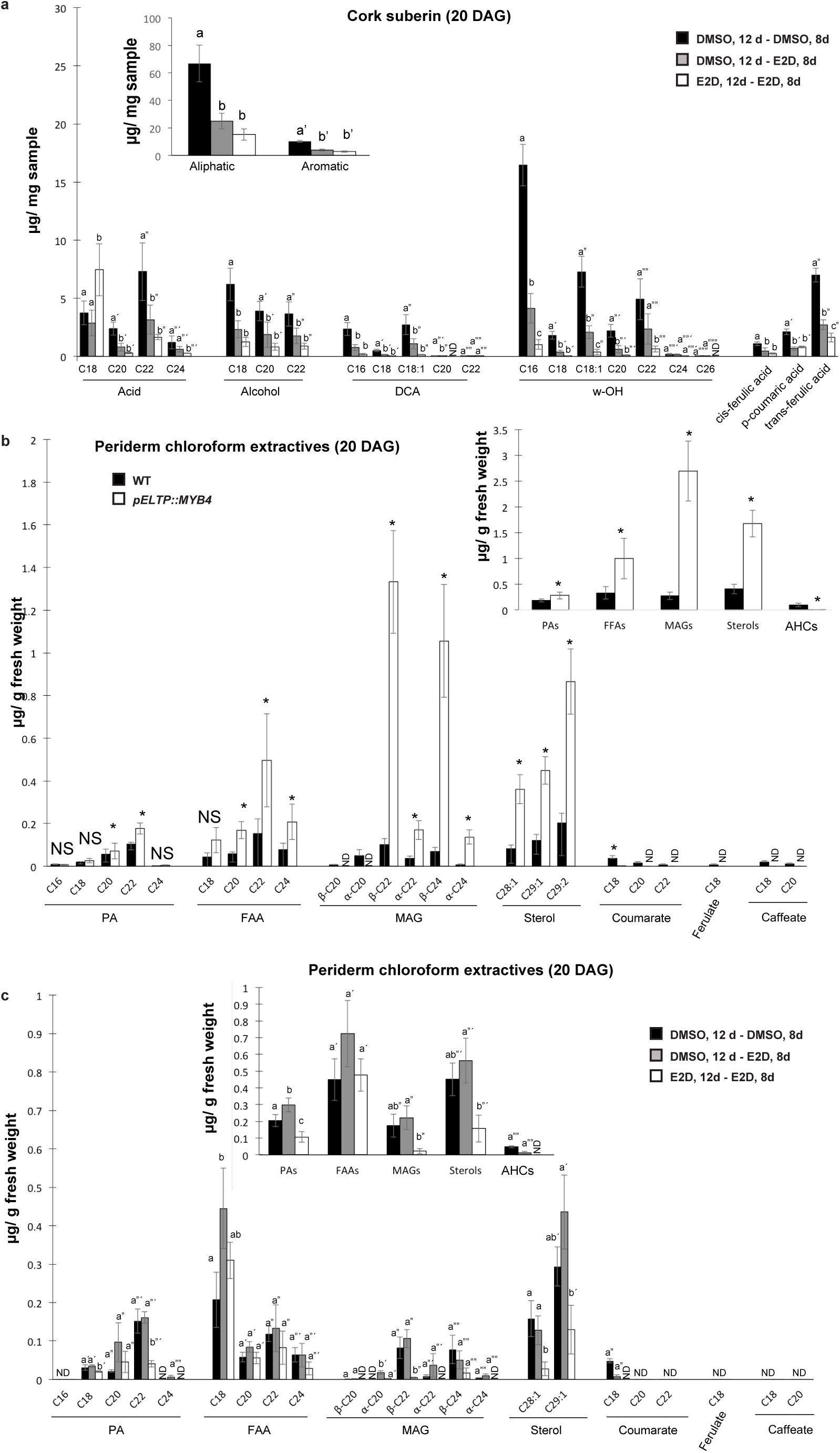
Suberin and Chloroform extractives quantification in the endodermis. a) Suberin quantification by Gas Chromatography coupled Mass Spectroscopy (GC-MS)-based analysis from the cork of p*ELTP*::X-VE>>MYB4-GFP expressing roots under different beta-estradiol (E2D) treatments (One-way ANOVA, CI95%, N=3-4). b) Chloroform extractives quantification by GC-MS-based analysis from the cork of 20-day-old WT and p*ELTP*::MYB4 expressing roots (N=3-4; T-Test *: P<0,05). c) Chloroform extractives quantification by GC-MS-based analysis from the cork of p*ELTP*::XVE>>MYB4-GFP expressing roots under different E2D treatments (One-way ANOVA, CI95%, N=3). For the comparison of the suberin fractions, each compound was normalized to the total aliphatic or aromatic respectively. Numbers refer to carbon chain lengths of aliphatic fatty acid derivatives. AHCs: Alkyl hydroxycinnamates, DCA: *α–ω*-dicarboxylic acid, FFAs: Free fatty acids, MAGs: Mono-acyl glycerol conjugates, ND: Not detected, NS: Not significant, PAs: Primary alcohols, All error bars are SD, letters refer to individual groups in a one-way ANOVA analysis.

**Supplementary Figure 8.**
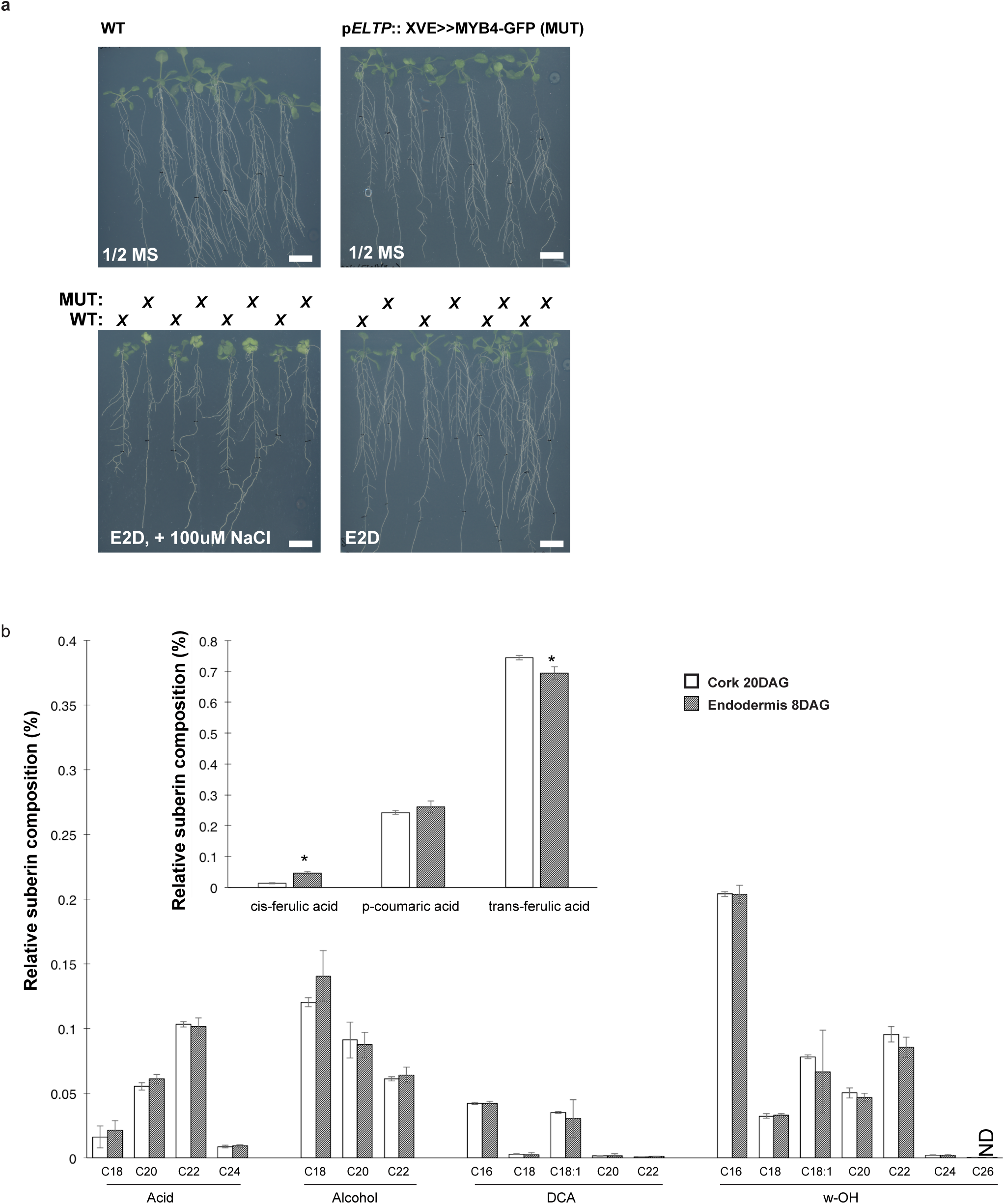
Phenotypes of p*ELTP*::XVE>>MYB4-GFP roots under stress. a) Images depicting p*ELTP*::XVE>> MYB4-GFP plants grown for 14 days under different regimes of the experiment shown in Figure 7d and 7e. Scale bars: 1cm. b) Relative composition of suberin of the cork of 20-day-old roots and the endodermis of 8-day-old roots. For the comparison, each compound was normalized to the total aliphatic or aromatic respectively. Numbers refer to carbon chain lengths of aliphatic fatty acid derivatives. DCA: *α–ω*-dicarboxylic acid, (N=3-5; T-Test *: P<0,05). All error bars are SD.

## Notes

### Competing Interest Statement

The authors have declared no competing interest.

